# Macrophage SLC9A1 Links Endocytic Trafficking to Innate Immune Activation in Myocardial Injury

**DOI:** 10.64898/2026.02.27.708643

**Authors:** Jinhua Wen, Pablo Parra, Yoshiharu Muto, Guo Chen, MariaSanta C Mangione, Xiang Luo, Dian J Cao

**Author notes:** Authors contributed equally (Jinhua Wen, Pablo Parra). To whom correspondence should be addressed: Dian Cao, MD, PhD, Division of Cardiology, UT Southwestern Medical Center, 6000 Harry Hines Blvd, Dallas, TX 75390-8573, Twitter: @DrDianCao.

## Abstract

Excessive innate immune activation drives adverse remodeling after myocardial infarction (MI), yet the upstream mechanisms by which macrophages sense ischemic danger signals remain poorly defined. Here we tested whether macropinocytosis functions as a mediator of post-ischemic inflammation and whether the Na⁺/H⁺ exchanger SLC9A1 links membrane ion transport to innate immune activation in the injured heart.

Macropinocytosis was robustly activated in infarct-associated macrophages, which are the predominant cell type with the macropinocytotic activity in the injured heart. Pharmacologic inhibition of macropinocytosis with 5-(N-ethyl-N-isopropyl)amiloride (EIPA) improved cardiac function and attenuated post-MI remodeling. EIPA also attenuated cardiac inflammatory responses induced by systemic lipopolysaccharide and Poly(I:C). To define macrophage-intrinsic mechanisms, we generated monocyte- and monocyte-derived macrophage–specific *Slc9a1* knockout mice. Genetic deletion of *Slc9a1* recapitulated the cardioprotective effects of EIPA and markedly suppressed interferon-stimulated gene programs in infarct-associated macrophages, as revealed by single-cell RNA sequencing. Mechanistically, SLC9A1 promoted endocytic uptake of Poly(I:C) acid and enhanced endosome-dependent inflammatory signaling.

Together, these findings identify macrophage macropinocytosis as a regulator of innate immune activation after MI and reveal SLC9A1 as a previously unrecognized link between membrane ion transport and inflammatory signaling in the injured heart. Targeting SLC9A1-dependent membrane trafficking pathways may therefore represent a strategy to limit maladaptive inflammation in ischemic heart disease.

## Introduction

Acute myocardial infarction (MI) triggers a robust inflammatory response that is essential for debris clearance, but, when excessive or prolonged, drives adverse cardiac remodeling and progression to heart failure. Innate immune activation is a central component of this response, with macrophages serving as key orchestrators of post-ischemic inflammation, tissue repair, and remodeling. Multiple pattern recognition receptors (PRRs) are activated in the infarcted myocardium by exposure to damage-associated molecular patterns (DAMPs) such as endogenous nucleic acids released from necrotic cardiomyocytes^1, 2^. However, despite extensive characterization of PRR signaling pathways, a key mechanistic question remains unclear: how extracellular danger signals in the MI environment gain intracellular access to engage endosomal innate immune sensors in post-infarction macrophages.

Macropinocytosis is a high-capacity, actin-dependent endocytic process by which cells internalize large volumes of extracellular fluid and soluble cargo, including proteins and nucleic acids^3–6^. Unlike receptor-mediated endocytosis, macropinocytosis is strongly induced by cellular stress, growth factor signaling, and tissue injury, conditions that are prominent within the ischemic myocardium. In myeloid cells, bulk endocytic uptake pathways such as macropinocytosis could deliver extracellular nucleic-acid cargo to endolysosomal compartments, enabling activation of intracellular PRRs and downstream inflammatory signaling^7^. The role macrophage macropinocytosis in inflammatory amplification following myocardial ischemia has not been examined.

Macrophages are highly plastic cells that continuously survey their microenvironment and dynamically adjust their functional states in response to local cues^8, 9^. To support this surveillance, macrophages internalize membrane equivalent to their entire cell surface every 30–35 minutes^10^, with macropinocytosis accounting for a substantial fraction of this uptake. Following MI, the necrotic myocardium releases abundant DAMPs, including nucleic acids and proteins, that are well suited for internalization via macropinocytosis. Once internalized, these danger signals may engage endosomal PRRs to initiate inflammatory signaling, while simultaneously providing metabolic substrates that support macrophage survival and mTORC1 activation in nutrient-restricted ischemic tissue^11^. Through these dual functions, macropinocytosis is positioned to coordinate inflammatory sensing and metabolic adaptation in infarct-associated macrophages.

Solute carrier family 9 member A1 (SLC9A1), also known as the Na⁺/H⁺ exchanger 1 (NHE1), is a plasma membrane antiporter that plays a fundamental role in intracellular pH and volume homeostasis. By extruding intracellular protons in exchange for extracellular sodium ions, SLC9A1 counteracts cytosolic acidification and facilitates osmotic swelling required for membrane expansion^12^. Beyond its canonical role in pH regulation, SLC9A1 regulates actin cytoskeletal organization, membrane ruffling, and endosomal dynamics^13, 14^, processes that are integral to macropinocytosis. Furthermore, Pharmacological inhibition of macropinocytosis has long been attributed to blockade of Na⁺/H⁺ exchangers, particularly SLC9A1/NHE1^15^, implicating SLC9A1 as a potential regulator of macropinocytosis-dependent immune sensing.

Despite studies of SLC9A1 in excitable tissues such as the heart and brain, its function in monocytes and macrophages remains poorly defined. Emerging evidence links SLC9A1 to macrophage-driven foam cell formation and atherogenesis^16^, yet historical studies of SLC9A1 in acute cardiovascular inflammation have largely focused on cardiomyocytes, leaving its role in macrophages unexplored. This gap is notable given that SLC9A1 is the predominant Na⁺/H⁺ exchanger isoform expressed in macrophages^17, 18^ and that therapeutic inhibition of this transporter regulated macrophage function in atherosclerosis^16^. Whether SLC9A1 regulates macrophage inflammatory responses during MI by controlling macropinocytosis and danger signal sensing has not been determined.

In this study, we investigated whether macropinocytosis functions as a critical regulator for innate immune activation following myocardial ischemia and whether SLC9A1 contributes to this process in monocytes and macrophages. We hypothesized that macropinocytosis-dependent uptake of danger signals amplifies post-MI inflammation and that SLC9A1 selectively enables this process by facilitating delivery of nucleic acid ligands to endosomal PRRs. To test this hypothesis, we first determined whether macropinocytic activity is engaged in macrophages within the infarcted myocardium and whether inhibition of this pathway confers protection from ischemic injury. To address the translational potential of targeting this process, we evaluated the effects of 5-(N-ethyl-N-isopropyl)amiloride (EIPA), an amiloride derivative with experimental and clinical use. We further extended these analyses to models that simulated systemic viral and bacterial challenge to determine whether pharmacologic inhibition with EIPA similarly modulates innate immune activation beyond myocardial injury.

To define the macrophage-intrinsic role of SLC9A1, we generated monocyte- and monocyte-derived macrophage (MDM)–specific deletion of *Slc9a1* and performed single-cell RNA sequencing (scRNA-seq) on infarct-associated MDMs. In parallel, we employed siRNA-mediated knockdown approaches and selective PRR ligands, Poly(I:C) for TLR3 and lipopolysaccharide (LPS) for TLR4, to delineate the mechanistic relationship between SLC9A1, macropinocytosis, and innate immune signaling.

## Materials and methods

### Animals

Animals were kept in a pathogen-free environment with free access to food and water. They were maintained on a 12-hour light/dark cycle from 6 am to 6 pm. All procedures were approved by the Institutional Animal Care and Use Committee at the University of Texas Southwestern Medical Center. The *Nhe1*^flox/flox^ mice were kindly provided by Dr. Dandan Sun from the University of Pittsburgh^19^. The Ccr2-CreER-GFP mice were from JAX (#:035229). The *Nhe1*^flox/flox^ and the *Ccr2-CreER-GFP* are cross-bred to generate the compound mice *Nhe1*^flox/flox^ and the *Ccr2-CreER-GFP.* To induce *Nhe1* deletion from monocytes, tamoxifen was administered via ip injection at 75 mg/kg for 3 days. Control cohorts, *Ccr2-CreER-GFP* and *Nhe1*^flox/flox^ received the same amount and duration of tamoxifen treatment. Baseline echocardiogram was performed before and after tamoxifen injection. All mice are from the C57BL/6J background.

### Reagents

The antibodies and the sources of these antbodies are listed in the following table. Reagents used are described in specific experiments.

| Antibody name listed by antigen | Cat# | Vendor |
| --- | --- | --- |
| P-S6 | 4858S | MyBiosource |
| S6 | MBS475585 | mybiosource |
| mTOR | MBS9131453 | Fitzgerald |
| GAPDH | 10R-G109A | Cell Signaling Technologies |
| Phospho-NFKB | 3033T | Cell Signaling Technologies |
| NFKB | 8242T | Fisher Scientific |
| CD68 | MCA1957GA | Cell Signaling Technologies |
| p-ERK | 9101S | Cell Signaling Technologies |
| ERK | 9107S | Cell Signaling Technologies |
| Toll-like Receptor 4 | 14358S | Cell Signaling Technologies |
| Phospho-Akt (Ser473) | 4060S | Cell Signaling Technologies |
| Phospho-Akt (Thr308) | 4056S | Cell Signaling Technologies |
| Phospho-p70 S6 Kinase (Thr389) | 9205S | Cell Signaling Technologies |
| AKT (pan) | 2920S | Cell Signaling Technologies |
| S6 | 2317S | Cell Signaling Technologies |
| Tuberin/TSC2 | 4308T | Cell Signaling Technologies |
| Phospho-Tuberin/TSC2 | 3617T | Cell Signaling Technologies |
| 4E-BP1 | 9644S | Cell Signaling Technologies |
| Phospho-4E-BP1 | 2855S | Life technologies |
| POLR3F | PA520589 | Cell Signaling Technologies |
| Phospho-PI3 Kinase p85 (Tyr458)/p55 (Tyr199) | 17366 | Cell Signaling Technologies |
| PI3 Kinase p85 | 4292 | Cell Signaling Technologies |
| IL-1β | #31202 | Proteintech |
| NHE1 | 29761-1-AP | Novus Biologicals |
| MyD88 | MAB3109-SP | Novus Biologicals |
| Phospho GSK3a/b (Ser21/9) | 9327S | Cell Signaling Tech. |
| GSK-3β | 610202 | BD |
| mTOR | 2972 | Cell Signaling Technologies |
| mTOR | 4517s | Cell Signaling Technologies |
| mTOR | 2983 | Cell Signaling Technologies |
| Phospho-mTOR | 2974 | Cell Signaling Technologies |

#### Primer sequences used for quantitative real-time PCR (qRT-PCR)

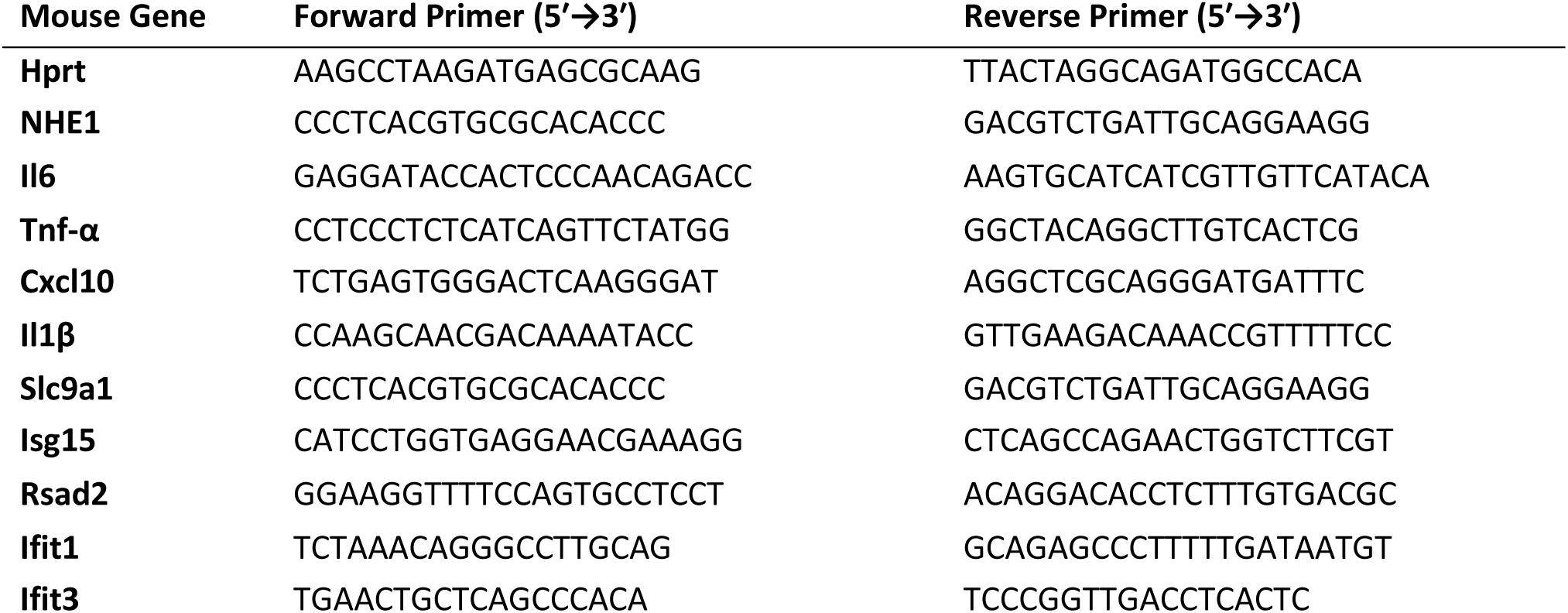

### Mouse myocardial infarction model

A murine model of myocardial infarction was employed involving permanent ligation of the left anterior descending coronary artery (LAD) as described previously^1, 11, 20^. Briefly, 10-16 weeks old mice were anesthetized with 2.4% isoflurane and positioned supine on a heating pad (37°C). Tracheal intubation was performed with a 19G stump needle, and animals ventilated with room air using a MiniVent mouse ventilator (Hugo Sachs Elektronik, stroke volume 250 μL, respiratory rate 210 breaths per minute). Following left thoracotomy between the fourth and fifth ribs, the LAD was visualized under a microscope and ligated with 6-0 Prolene suture. The suture was placed at the LAD segment corresponding to 1.5 mm to 2 mm below the lower edge of the left atrium. Regional ischemia was confirmed by visual inspection of discoloration of myocardium distal to the occlusion under a dissecting microscope (Leica). Sham-operated animals underwent the same procedure without occlusion of the LAD. After LAD ligation, hearts were collected at 24-, 48-, 72-hour, and 1-week. Left ventricles were separated into remote and infarct-at-risk zones for downstream analysis.

### Bone marrow monocytes isolation

Mice were euthanized and the surface is sprayed with the 70% ethanol. The soft tissues were removed from the femurs with a sterile scalpel and the clean bones (n = 4) were transferred into a petri dish on ice. Both ends of the long bone (epiphysis) of the femur were cut to expose the bone marrow (BM). The PBS was used to flush out the bone marrow and collected in a 15 mL tube. The BM cell suspension was centrifuged at 300 g for 5 minutes, and the pellet was resuspended in cell sorting buffer containing 2% FBS and 1mM EDTA and filtered using a 40µm cell strainer immediately before isolation. We use Fluorescence-activated cell sorting (FACS) method to purify neutrophils. We label the BM single cell suspension with CD45, CD11b, Ly6G, CCR2, and Ly6C. The CD45^+^CD11b^+^Ly6G^-^Ly6C^hi^ cell population (monocytes) is sorted and collected for downstream analysis. In mice with CCR2-Cre-GFP, the endogeneous GFP is used as a parameter to sort monocytes (CD45^+^CD11b^+^Ly6G^-^CFP^+^).

### Cardiac macrophage isolation

Cardiac immune cell isolation as performed based on our published protocol ^1, 11, 20^. Briefly, the MI tissue was separated from the remote myocardium and rinsed in 4°C PBS. The tissue was extensively rinsed to remove circulating blood, then minced with surgical scissors. The minced tissues were then incubated at 37°C for 45 minutes in a Thermomixer (Eppendorf) with shaking set at 120rpm in the digestion buffer that contained the following enzymes (all from Sigma, St. Louis, MI): collagenase I (cat. no. SCR103,) 450 unit/mL, hyaluronidase (cat. No. H3884) 60 unit/ml, DNase (cat. 900 3-98-9) 60 unit/ml. The tissues were then triturated and passed through a 40 μm strainer. Cells were then washed in 50 ml 4°C PBS containing 1% Fetal Calf Serum (FCS) by centrifugation for 10 minutes at 400 g. The resulting cell pellet was resuspended in cell sorting buffer before FACS, flow cytometry, or immunomagnetic isolation using the isolation kit from Miltenyi (cat.130 -120-337).

### Flow cytometry

The single cell suspension from the infarct tissue, peripheral blood (PB), and bone marrow (BM) was obtained and the cell concentration was adjusted to 1×10^6^/100 μL with sorting buffer. The cells were incubated first with purified anti-mouse CD16/32 (FcBlock) for 10 minutes at 4 °C followed by adding antibodies to the concentration specified in the table. The cells were incubated with antibodies at 4 °C for 15 minutes. After incubation, the cells were pelleted by centrifugation at 500g for 5 minutes and washed twice in the sorting buffer. The labeled cells were analyzed by flow cytometry. Gating strategies are described in the Results section. Mean fluorescence intensity was calculated as the median of the indicated fluorescent parameter using FlowJo software. Cell sorting was performed on an Aria III Fusion (BD Biosciences). No sodium azide in the FACS buffer for sorting experiments. Antibodies used for flow cytometry: from Biolegend: CD45-PE/Cy7 (cat# 103114), CD11b-PB (cat# 101223), F4/80-PE (cat#123109), Ly-6G-APC (cat# 127613), Ly-6C-BV711 (cat# 128037), CD206-PerCP/Cy5.5 (cat# 141715), CD64-FITC (cat# 164405), CCR2-BV421 (cat# 150605).

### Preparation of myocardial DAMP (mDAMP)

The remote heart tissue from mice, 3 or 4 days post-myocardial infarction (MI) surgery, was collected for mitochondrial isolation. Briefly, approximately 250 mg of heart tissue was cut into 1 –2 mm pieces and rinsed in cold PBS to remove any residual blood. The tissue was then transferred to a homogenizer containing 1 mL of cold mitochondrial isolation buffer and homogenized on ice. The resulting supernatant was collected, and the pellet was discarded by centrifugation at 1,000 g for 5 minutes at 4°C. The supernatant was further centrifuged at 12,000 g for 10 minutes at 4°C. The resulting pellet was used as the mitochondrial fraction and subjected to freeze-thaw cycles on dry ice (5 times). mDAMPs were generated by centrifugation after the freeze-thaw cycles at 12,000 g for 10 minutes and stored at -80°C.

### Extracellular fluid extraction from myocardial infarction tissue

Infarcted heart tissue collected 2-3 days after the MI surgery was used to extract extracellular fluid by centrifugation^21^. Briefly, heart tissue was washed in cold PBS and sectioned into small pieces. 200mg of samples were transferred to an insert with a nylon mesh (20 μm opening, 55 μm thickness, Spetrum Labs) placed on top of a mini spin column (Qiagen), from which the bottom filter was removed. The tubes were centrifuged at 400 g for 15 minutes at 4°C, and then the inserts were removed. The fluid was centrifuged for another 5min art 400 g at 4°C. The fluid from 3 samples was pooled and spun for 10min at 4000 g at 4°C. The supernatant was stored at -80°C upon proteomic analysis.

### Culture to develop Bone marrow-derived macrophages (BMDM)

C57BL6/J female mice, aged 6–8 weeks and weighing 20–25 g, were used to isolate bone marrow cells from their two femurs and two tibias^22^. Bone marrow cells from one mouse were incubated in 10 mL RPMI-1640 (Bio Life Sciences) supplemented with 10% Fetal Bovine Serum (FBS, Sigma Aldrich), and then filtered through a 40 μm Fisherbrand™ Sterile Cell Strainer (Fisher Scientific). The cells were collected by centrifugation at 300 g for 5 minutes at 4°C and resuspended in RPMI -1640 with 10% FBS and 20 ng/mL recombinant mouse M-CSF (R&D systems). The cells were cultured in two 10-cm Petri dishes at 37°C. After 3-4 days, the culture medium was replaced, and the cells were cultured until day 6-7. BMDMs were then used for characterization and stimulation experiments.

### BMDM Stimulation with mDAMP, EIPA, and NPS

For stimulation, 3×10⁵ BMDMs per well were plated in a 12-well plate and cultured overnight at 37°C. The cells were then starved in FBS-free RPMI-1640 for 3 hours. The cells were treated with 100 μg/mL MDAMP protein with or without 12.5 μM 5-n-ethyl-n-isopropyl amiloride (EIPA, Fisher Scientific), 3 μM NPS-2143 (Fisher Scientific), and incubated at 37°C for 3 hours. After the treatment, the cells were collected for protein and RNA extraction.

### siRNA-mediated *Slc9a1* silencing

*Slc9a1* in BMDMs were silenced using small interfering RNA (siRNA) (Life Technology Corporation). The sequence sense 5′-CCACAAUUUGACCAACUUAtt-3′ and antisense 5′-UAAGUUGGUCAAAUUGUGGtc-3′ was used to inhibit *Slc9a1* expression and a non-specific sequence sense 5′-UUCUCCGAACGUGUCACGUdTdT-3′and antisense 5′-ACGUGACACGUUCGGAGAAdTdT-3′ were used as a negative control. Cells were transfected with 40 nM *Slc9a1*-targeting or non-targeting control siRNA using Lipofectamine RNAiMAX according to the manufacturer’s instructions. Briefly, Cells were seeded at 3 × 10⁵ cells per well in 12-well plates 18–24 hours prior to transfection. For each well, siRNA and Lipofectamine RNAiMAX were diluted separately in Opti-MEM and then combined, followed by additional incubation for 5-10 minutes at room temperature to allow complex formation. Complexes were added dropwise to cells with serum-free medium. Medium was replaced to complete medium after 24 hours. Cells were harvested or treated for downstream analysis at 72-96 hours of transfection.

### Poly(I:C) and LPS stimulation

High-molecular weight polyinosinic–polycytidylic acid (Poly(I:C) (InvivoGen) and Lipopolysaccharides (LPS) (Sigma) were reconstituted in sterile endotoxin-free water at 1 mg/mL. BMDMs were maintained at 37 °C in a humidified incubator containing 5% CO₂ in complete growth medium with 20 ng/ml of M-CSF. Cells were seeded 18–24 h prior to stimulation to achieve 70–80% confluence, and the medium was replaced with serum-free medium at the time of treatment. Poly(I:C) or LPS was added directly to the culture medium at a final concentration of 5 μg/ml or 10 to 100 ng/mL, respectively. Cells were treated for 4 h and collected for downstream assays.

### Rhodamine-Poly(I:C) uptake assay

High molecular weight Rhodamine-Poly(I:C) (InvivoGen) was used to assess cellular uptake. BMDMs were seeded at 1-2 × 10⁵ cells per well in 24-well plates with coverslips and cultured overnight in complete medium at 37°C in 5% CO₂ to allow adherence. Prior to the uptake assay, cells were incubated in serum-free medium to minimize interference from serum components. After incubated with 1 μg/ml of Rhoamine-Poly(I:C) (InvivoGen) at 37°C for 1 h or 4 h, the cells were washed three times with ice-cold PBS to remove unbound ligand and fixed in 2% paraformaldehyde for 15 minutes at room temperature. Coverslips were mounted with an antifade reagent with DAPI. Images were acquired under identical confocal microscopy settings for all groups.

### Purification of bone marrow monocytes by FACS

Bone marrow cells were isolated from 8-12-week-old mice. Femurs and tibias were harvested under sterile conditions. Bone marrow was flushed using a 27-gauge needle with cold sorting buffer (PBS + 0.5 % BSA + 1 mM EDTA). Cell suspensions were passed through a 40-µm cell strainer to obtain single-cell suspensions. Red blood cells were lysed with ACK lysis buffer for 2 minutes at room temperature, then washed with sorting buffer. For surface marker staining, cells were incubated with normal Rat serum for 10 minutes at room temperature to prevent nonspecific binding. Cells were then stained for 30 minutes at 4°C under light-free conditions with fluorochrome-conjugated antibodies diluted in sorting buffer. A typical monocyte panel (CD45 ^+^CD11b^+^Ly6G^-^GFP^+^) was used to select the monocytes from CCR2-CreER-GFP and *Slc9a1*^flox/flox^;CCR2-Cre-GFP mice. To isolate r monocytes from WT or the *Slc9a1*^f/f^ mice, we gated the population as CD45^+^CD11b^+^Ly6G^-^Ly6C^hi^CCR2^+^. After staining, cells were washed and resuspended in sorting buffer for cell sorting on the FACS system. The sorted cells were aliquoted for further assays.

### In vivo Poly(I:C) and LPS challenge model

Poly(I:C) and LPS were prepared fresh in sterile saline. EIPA was dissolved in DMSO. Wild-type mice were randomly assigned to vehicle or EIPA controls, Poly(I:C) alone, LPS alone, EIPA + Poly(I:C) and EIPA + LPS groups. EIPA was administered subcutaneously (s.c) at 15 mg/kg body weight/day at 1 day and 2 hours prior to Poly(I:C) and LPS challenge. To induce the systemic inflammatory responses, Poly(I:C) or LPS was administered via intraperitoneal (i.p.) injection at 4 mg/kg body weight or 1 mg/kg body weight, respectively. Tissues (spleen, bone marrow, heart) were collected at 4 hours post-injection to perform mRNA analysis by RT-qPCR and protein production detection by western blot assay. In separate cohorts, WT mice were challenged with 10 mg/kg LPS after receiving DMSO or EIPA injection (15 mg/kg) daily for three days. The LPS challenge was administered 2 h after the last dose of DMSO or EIPA. Echocardiograms were performed at 6 h and 24 h after LPS. Mice were sacrificed, and tissues (cardiac, BM, and spleen) were collected at 24 h post-LPS injection.

#### Immunoblot

Cells were harvested, lysed in RIPA buffer (Fisher Scientific) supplemented with protease and phosphatase inhibitors (Protease and Phosphatase Inhibitor Mini Tablets, EDTA-free, Fisher Scientific) and then incubated on ice for 15 minutes. The lysate was clarified by centrifugation at 12,000 x g for 10 minutes at 4°C. The protein-containing supernatant was collected, and protein concentration was determined using a BCA protein assay. The protein sample was diluted with 5X loading buffer and heated at 95°C for 5 minutes to denature the proteins. Protein lysates were separated by SDS-PAGE, transferred to Hybond-C nitrocellulose membranes, and subjected to immunoblot analysis. The membranes were incubated with the primary antibody while gently shaking overnight at 4°C. After incubation, the membranes were washed three times for 5 minutes each with TBS-T (tris-buffered saline with 0.1% Tween-20). Next, the membranes were incubated with IRDye 800CW or 700CW secondary antibodies (LI-COR) for 1 hour at room temperature with gentle shaking. The membranes were washed again three times for 5 minutes each with TBS-T. Protein bands were visualized using a LI-COR Odyssey Imaging System, and band intensity was quantified using the Image Studio software.

#### RNA extraction and RT-qPCR

Total RNA was extracted from BMDM cells using the Quick-RNA™ Microprep Kit (Zymo Research). Briefly, 1 volume of 95–100% ethanol was added to 1 volume of the sample lysed in RNA Lysis Buffer (1:1). The mixture was then transferred to a Zymo-Spin™ IC Column placed in a Collection Tube and centrifuged. After DNase I treatment, the RNA was washed and eluted from the column using DNase/RNase-Free Water. The RNA was subsequently used in a reverse transcription reaction with PrimeScript™ RT Master Mix (TAKARA) and then analyzed by qPCR using iTaq Universal SYBR Green Supermix (Bio-Rad).

### Macropinocytosis assay

BMDM cells were incubated with a 500 µg/mL solution of dextran 70,000 MW (Oregon Green 488, anionic, lysine-fixable) for 40 minutes. The cells were then fixed with 4% PFA for 15 minutes and washed three times with cold PBS. The cell nuclei were stained using Vybrant® DyeCycle™ Violet stain (Invitrogen).

For in vivo dextran uptake investigation, on day 3 after underwent myocardial infarction (MI) surgery, the mice were anesthetized, and the hearts were carefully exposed. A 1 mg/mL solution of dextran 70,000 MW was locally perfused on the surface of the hearts for 30 minutes. The mice were subsequently euthanized, and the hearts were excised immediately above the suture line, then rinsed in ice-cold PBS. The tissues were fixed in 4% PFA at 4°C with gentle shaking for 3 hours. To cryopreserve the heart tissue, it was embedded in OCT compound, snap-frozen in liquid nitrogen, and stored at -80°C.

#### Histology, Immunohistochemistry, and Immunocytochemistry

Following euthanasia via cervical dislocation, the animals’ hearts were carefully rinsed in PBS and fixed in 4% paraformaldehyde (PFA) at room temperature with gentle agitation. The tissues were then processed for paraffin embedding or Cryo-section using standard protocols. For immunostaining, 5-µm coronal sections were prepared to obtain a four-chamber view of the heart. The sections were permeabilized with 0.25% Triton X-100 in PBS and then blocked with 5% BSA for 1 hour at room temperature. The primary antibodies were applied and incubated overnight at 4°C. After washing three times with PBS, the sections were incubated with the secondary antibody for 1 hour at room temperature. The sections were then mounted with ProLong™ Gold Antifade Mountant containing DAPI to stain the nucleus. Hematoxylin and eosin (H&E) staining was performed according to standard protocols.

Cultured BMDM cells were washed three times with PBS, then fixed in 4% PFA for 15 minutes. Cells were permeabilized with 0.1% Triton X-100 for 15 minutes and subsequently blocked in PBS containing 5% BSA for 1 hour. The cells were then incubated with primary antibody. Donkey- or goat-derived secondary antibodies conjugated to Alexa 488 or 546 were used to visualize the fluorescence signals. ProLong™ Gold Antifade Mountant containing DAPI was used to stain the nucleus.

### Confocal images were analyzed using ImageJ (NIH)

For quantification of Rhodamine–Poly(I:C) uptake or EEA1 staining, images from all experimental groups were acquired under identical microscope settings (laser power, gain, and exposure) to allow direct comparison. Raw images were first converted to grayscale and background subtraction was applied uniformly across all images. A fluorescence intensity threshold was then determined empirically from control samples and applied uniformly to all images within the experiment to define positive signal. Cells exceeding this threshold were scored as positive. For each field, the percentage of positive cells was calculated relative to the total number of cells identified by nuclear staining or brightfield morphology. In addition, fluorescence intensity measurements were extracted using the ImageJ “Measure” function, including mean fluorescence intensity, integrated density, and raw intensity per cell or region of interest. All analyses were performed using the same threshold parameters across experimental groups to ensure unbiased comparison.

### Single-cell RNA sequencing of the macrophages purified from the MI tissue

Single-cell suspensions from infarcted cardiac tissue were generated as previously described ^1, 11, 20^. Immune cells were first enriched by positive selection using a CD45 MicroBeads kit according to the manufacturer’s instructions (130-052-301, Miltenyi Biotec). The enriched cell fraction was subsequently stained with antibodies against CD45, CD11b, and CD3, and CD45⁺CD11b⁺CD3⁻ cells were isolated by fluorescence-activated cell sorting. Sorted cells were assessed for viability (>85%) and immediately processed for single-cell RNA sequencing. Single-cell libraries were prepared using the Chromium Single Cell 3′ Gene Expression platform (10x Genomics) following the manufacturer’s protocol. Briefly, single cells were partitioned into nanoliter-scale Gel Bead-in-Emulsions (GEMs), where cell lysis and barcoded reverse transcription occurred, enabling the incorporation of unique cell barcodes and unique molecular identifiers (UMIs) into cDNA. Following GEM recovery and cDNA amplification, sequencing libraries were constructed and sequenced on an Illumina platform. Raw sequencing data were processed using the Cell Ranger pipeline (10x Genomics) to generate gene-by-cell expression matrices for downstream analysis.

### Bone marrow monocyte and cardiac macrophage isolation and bulk RNA sequencing

A detailed protocol for bulk RNA sequencing was described in our prior work^20^.

### Echocardiography

Mouse echocardiograms were recorded on conscious, gently restrained mice using a Vevo 2100 system and an 18-MHz linear probe as described^1, 11, 20^. A short-axis view of the LV at the level of the papillary muscles was obtained, and M-mode recordings were obtained from this view. Left ventricular internal diameter at end-diastole (LVIDd) and end-systole (LVIDs) were measured from M-mode recordings. Fractional shortening was calculated as (LVIDd − LVIDs)/LVIDd (%).

### Statistical analysis

Findings are expressed as mean ± SD. The data were analyzed using statistical software (GraphPad Prism, version 7.01; GraphPad Software, San Diego, CA). An unpaired Student t test was performed to analyze 2 independent groups. One-way ANOVA, followed by Tukey’s post hoc test, was used for pairwise comparisons. In representative datasets, we have also employed nonparametric tests (Wilcoxon rank sum test, Wilcoxon two-sample test, Kruskal-Wallis test). A value of P< 0.05 was considered statistically significant and numerical P values are displayed on results.

## Results

### Macropinocytosis Is Activated in Macrophages in the Infarct Tissue and Pharmacologic Inhibition Attenuates Post-Ischemic Inflammation and Myocardial Injury

By internalizing large quantities of soluble content from the environment, macropinocytosis could internalize both DAMPs and nutrients from the ischemic myocardium that promote macrophage activation and survival. We first determined if MI macrophages have active macropinocytosis. Macropinocytosis is conventionally quantified using high–molecular weight tracers, such as ∼70-kDa dextran or comparably sized fluorescent beads, which are preferentially internalized through fluid-phase bulk uptake pathways. Therefore, we performed LAD ligation and then isolated immune cells from the infarct tissue at four <u>d</u>ays <u>p</u>ost injury (4-dpi). The immune cells were incubated with FITC-Dextran 70 (FITC-D70) for thirty minutes. Figure 1A depicts the gating strategy used to identify the MI macrophages (CD45^+^CD11b^+^Ly6G^-^), neutrophils (CD45^+^CD11b^+^Ly6G^+^), and lymphocytes (CD45^+^CD11b^-^CD3^+^). As demonstrated in Figure 1B, macrophages were the main cells that engulf FITC-D70. Neutrophils, which are also phagocytes, engulfed D70 to a much lesser extent, whereas lymphocytes showed negligible FITC-D70 uptake. Next, we devised a model for examining macropinocytosis in vivo. DQ-Red BSA (67 kDa), a self-quenched fluorescent substrate that emits red fluorescence upon intracellular proteolytic processing, is commonly used as a functional readout of macropinocytic uptake^23^. We infused DQ-Red BSA into the infarct tissue for 40 minutes, followed by immediate removal of the heart for cryosection. Figure 1C shows that degraded (i.e. fluorescent) DQ-Red BSA primarily colocalized with CD68^+^ macrophages in the infarct region. These data suggest that MI macrophages are undergoing macropinocytosis and are the major cells in the ischemic environment with this activity.

**Figure 1.**
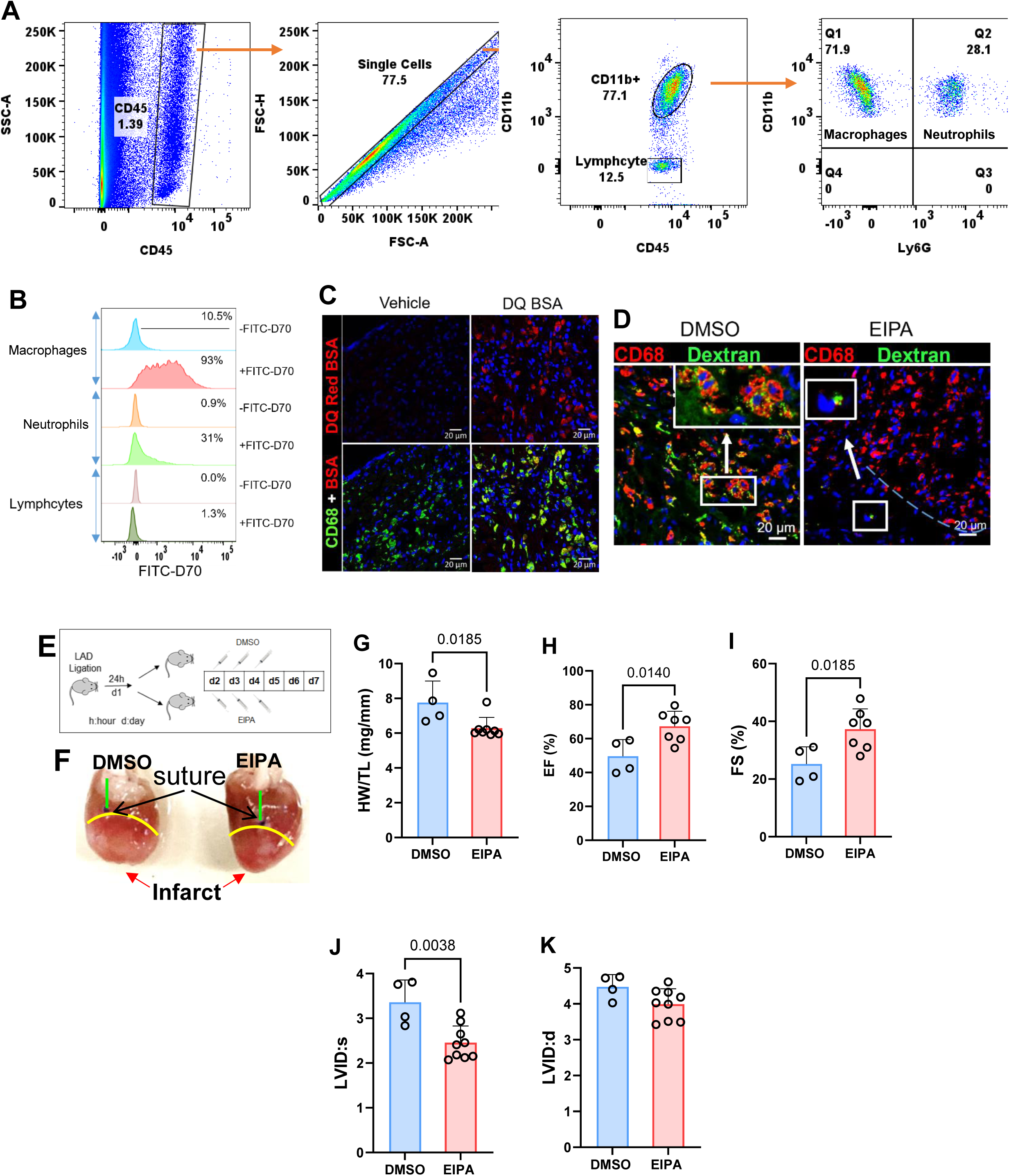
Macropinocytosis is activated in infarct-associated macrophages, and pharmacologic inhibition with EIPA attenuates ischemic injury. Myocardial infarction was induced by permanent ligation of the LAD. A. Experimental strategy for detecting macropinocytosis in infarct tissue. Single-cell suspensions generated from infarct tissue 4dpi were incubated with FITC–dextran (70 kDa; D70) for 40 minutes and stained with CD45, CD11b, and Ly6G. Macrophages (CD45⁺CD11b⁺Ly6G⁻), neutrophils (CD45⁺CD11b⁺Ly6G⁺), and non-myeloid cells (mostly lymphocytes) (CD45⁺CD11b⁻Ly6G⁻) were gated. B. Percentage of cells positive for FITC–D70 uptake in the indicated populations. C. DQ-BSA uptake in infarct tissue. DQ-BSA was delivered into the myocardium at 4dpi, and tissue sections were stained with CD68. DQ-BSA fluorescence (red) colocalized with CD68⁺ macrophages. D. FITC–D70 uptake in infarct tissue from mice treated with vehicle (DMSO) or EIPA (12.5 mg/kg). Heart sections were stained for CD68. E. Experimental scheme for EIPA administration following MI. F. Representative heart morphology from vehicle- and EIPA-treated mice. G-K. Quantification of post-MI remodeling and cardiac function, including heart weight/tibia length ratio (HW/TL), ejection fraction (EF), fractional shortening (FS), and left ventricular end-systolic and end-diastolic dimensions. Student’s t-test (G-K).

Next, we tested how inhibiting macropinocytosis affects outcomes of MI. We injected EIPA (5-(N-ethyl-N-isopropyl)amiloride), a widely used agent to suppress macropinocytosis, or vehicle control before locally infusing the FITC-D70 into the infarct heart for 40 minutes while keeping the mice on a respirator. After harvesting and preserving the heart, cardiac tissue sections were labeled with macrophage marker CD68. As demonstrated in Figure 1D, FITC-D70 was readily detectable and all FITC-D70 signals colocalized with CD68^+^ macrophages (inset in Figure 1D). In mice that were pretreated with EIPA, the uptake of D70 was diminished (Figure 1D), suggesting EIPA inhibited macropinocytosis in MI macrophages.

To determine if inhibiting macropinocytosis by EIPA affects myocardial repair, we generated MI and then at 24 hours post-surgery administered vehicle control (DMSO) or EIPA at 12.5 mg/kg/day for a total of three days (Figure 1E). Figure 1F are images of hearts collected from DMSO and EIPA-treated mice at 7-dpi. DMSO-treated hearts showed loss of the conical shape of the LV apex, which is consistent with remodeling. EIPA treatment reduced the gravimetric indexes of remodeling including heart weight (HW) to tibia length (TL) ratio (Figure 1G), improved EF and FS (Figures 1H, 1I), and decreased LV systolic dimension (Figure 1J).

In aggregate, these data suggest that macropinocytosis is a robustly active process in MI macrophages and that blocking macropinocytosis protects the heart from ischemic injury.

### Myocardial DAMP Promotes Macropinocytosis and Is a Strong Activator of the TLR Pathway

Next, we designed an in vitro system to assess how effectively EIPA inhibits macropinocytosis in a simulated infarct environment. We extracted soluble content using enriched mitochondria preparation from the heart; we call this preparation “<u>m</u>yocardium DAMP (mDAMP)”. To establish the physiological relevance of this approach, we isolated interstitial fluid from infarcted myocardium (i-IF) at 48–72 hours after myocardial infarction, a time window during which macrophages are actively exposed to necrosis-derived danger signals in vivo. The DAMP components present in i-IF therefore represent the endogenous molecular environment encountered by macrophages within the infarct tissue. We then performed comparative proteomic analyses of mDAMP and i-IF. As shown in Figures 2A and 2B, both preparations contained a broad array of known DAMP molecules. Notably, more than 90% of annotated DAMP proteins^24^, including established cardiac DAMPs such as myosin^25^ and cytochrome C^26^, were shared between mDAMP and i-IF, although the relative abundance of individual proteins varied between the two preparations. The strong overlap indicates that mDAMP closely recapitulated the endogenous DAMP milieu of the ischemic myocardium and can be used as a biologically relevant stimulus for interrogating macrophage signaling pathways activated in vivo following myocardial injury.

**Figure 2.**
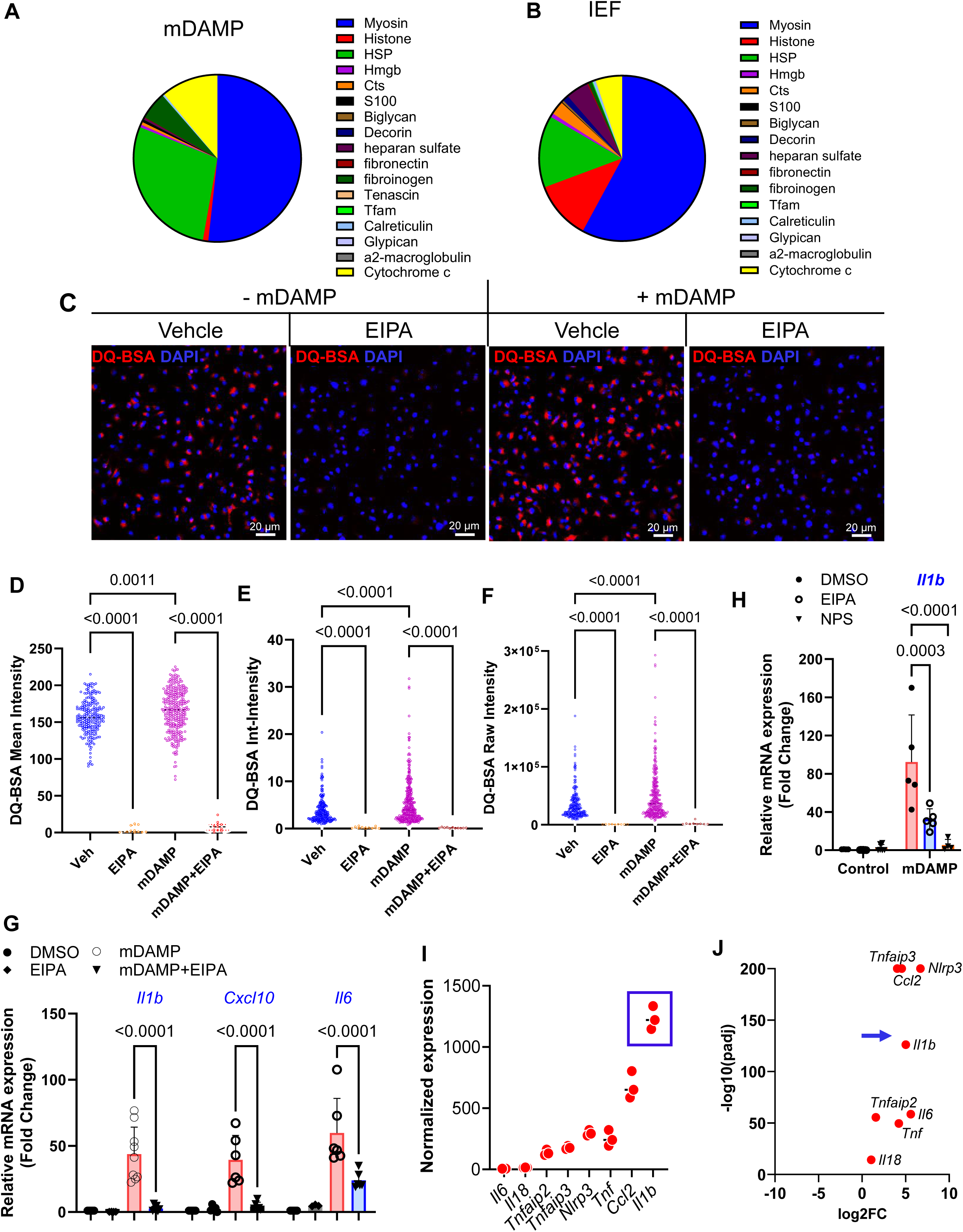
Inhibition of macropinocytosis suppresses mDAMP-induced macrophage activation. A-B. Proteomic analysis of mDAMP generated from enriched myocardial mitochondria (A) and the extracellular fluid isolated from infarct tissue (IEF) (B). C. BMDMs were stimulated with mDAMP in the presence or absence of EIPA (10 μM). The uptake of DQ-BSA was used to gauge macropinocytosis. Representative confocal images show intracellular DQ-BSA fluorescence (red). D-F. Quantification of DQ-BSA fluorescence using ImageJ, including mean intensity (D), integrated intensity (E), and raw fluorescence intensity (F). EIPA markedly reduced DQ-BSA uptake. G. BMDMs were stimulated with mDAMP in the presence or absence of EIPA (10 μM) for 4 h, followed by RNA extraction and RT-qPCR analysis. Expression of *Il1b*, *Cxcl10*, and *Il6* induced by mDAMP was reduced by EIPA. H. BMDMs were stimulated with mDAMP as in (G) and treated with EIPA or the structurally distinct macropinocytosis inhibitor NPS (3 μM). NPS markedly suppressed mDAMP-induced *Il1b* expression. I. Bulk RNA sequencing of macrophages isolated from infarct tissue by fluorescence-activated cell sorting (FACS). *Il1b* was among the most highly expressed inflammatory cytokines. J. Bulk RNA sequencing comparing bone marrow monocytes and infarct-associated macrophages purified by FACS. *Il1b* was among the most strongly upregulated cytokines in infarct macrophages. 2-way Anova

Next, we asked if mDAMP promotes macropinocytosis. We stimulated bone marrow-derived macrophages (BMDMs) with mDAMP and DQ-BSA for 40 minutes in the presence or absence of EIPA and quantified the uptake using fluorescent microscopy. As demonstrated in Figure 2C and the summarized data in Figures 2D, 2E, and 2F, mDAMP-treated cells had more uptake of DQ-BSA and while blocked both constitutive/baseline and mDAMP-stimulated DQ-BSA uptake, indicating it effectively suppressed macropinocytosis in macrophages.

Next, we sought to leverage the mDAMP to define the signaling pathways most likely regulated by macropinocytosis in the ischemic environment. Treatment of BMDMs with mDAMP induced rapid phosphorylation and activation of NF-κB (<u>Suppl</u>ementary (Suppl.) Figures 1A and 1B) and immediate and sustained activation of ERK (Suppl. Figures 1C and 1D). Additionally, mDAMP robustly activated mTORC1 signaling, as evidenced by increased phosphorylation of S6 (Suppl. Figure 1E) and hyperphosphorylation of 4EBP1 (Suppl. Figure 1F), which relieves translational repression and reflects enhanced mTORC1 activity. Quantification of mTORC1 activation is summarized in Suppl. Figures 1G and 1H. Torin 1, a potent mTOR inhibitor, was included as a control. Notably, similar to ERK, mDAMP-induced mTORC1 activation persisted throughout the experimental time course (Suppl. Figure 1I). The coordinated activation of NF-κB, ERK, and mTORC1 represents a well-established signaling signature of Toll-like receptor (TLR) engagement, particularly downstream of TLR4 ligands^27^. These findings suggest that mDAMP activates macrophages through TLR-mediated inflammatory signaling and that inhibiting macropinocytosis may disrupt inflammation downstream of TLR activation.

### Pharmacologic inhibition of macropinocytosis suppresses mDAMP-induced inflammatory response in macrophages

The compositional similarity between mDAMP and interstitial fluid from the infarct, the activation by mDAMP of traditional TLR-activated pathways, and the way mDAMP increased macropinocytosis makes mDAMP a useful tool for gauging the consequences of macropinocytosis for inflammatory activation after MI. We first determined whether mDAMP directly induced inflammatory cytokine expression in bone marrow–derived macrophages (BMDMs) and whether macropinocytosis plays a role therein. We examined a broad panel of inflammatory mediators, including chemokines of the CCL family (*Ccl2*, *Ccl4*, *Ccl5*), CXCL family (*Cxcl1*, *Cxcl2*, *Cxcl10*), interferons and interferon-stimulated genes (*Ifnb*, *Ifng*, *Irf7*, *Ifit1*, *Ifit3*, *Isg15*, and *Rsad2*), as well as proinflammatory cytokines (*Il1a*, *Il1b*, *Il18*, *Tnfa*, and *Il6*). Among these targets, *Il1b*, *Cxcl10*, and *Il6* were consistently and robustly induced by mDAMP stimulation (Figure 2G), a cytokine profile characteristic of TLR-driven inflammatory activation, particularly downstream of TLR4.

Importantly, treatment with EIPA almost completely abrogated mDAMP-induced *Il1b* and *Cxcl10* expression and significantly reduced *Il6* induction (Figure 2G). To exclude compound-specific effects, we next tested NPS, a structurally distinct macropinocytosis inhibitor. NPS similarly abolished mDAMP-induced *Il1b* expression (Figure 2H), supporting that macropinocytosis has a necessary role in mDAMP-driven macrophage activation and likely the ischemic tissue. As additional control, we also treated BMDMs with the mitochondria-enriched preparation, the precursor we used to extract mDAMP. In contrast to mDAMP, the precursor failed to induce inflammatory cytokine expression, including *Il1b* and *Il6* (Suppl. Figures 2A-2B), indicating that macrophage activation requires DAMP released from damaged mitochondria/myocardium. Furthermore, the precursor did not induce protein expression NLRP3, iNOS, and IL1B (Suppl. Figure 2C). Taken together, these data demonstrate that inhibiting macropinocytosis efficiently suppressed mDAMP-triggered inflammatory response in macrophages.

To link these in vitro observations to the in vivo disease context, we performed RNA sequencing analysis of purified cardiac macrophages isolated from infarcted myocardium and compared them with bone marrow monocytes, the precursors of infarct macrophages, which are named as monocyte-derived macrophages (MDM). Because monocytes undergo substantial functional reprogramming upon recruitment to injured tissue, comparing monocytes with MDMs enables identification of pathways specifically induced by the ischemic environment and provides insight into the molecular programs that govern macrophage activation within the infarcted heart. Strikingly, *Il1b*, a key inflammatory cytokine classically induced downstream of TLR4 signaling, was the most abundantly expressed gene in infarct cardiac macrophages. Among major inflammatory mediators implicated in post-MI inflammation, including TNF family members (*Tnfaip2*, *Tnfaip3*, *Tnf*)*, Ccl2*, *Il6*, *Il18,* and *Nlrp3*, *Il1b* was the highest expressed cytokine (Figure 2I, blue box). Moreover, *Il1b* expression was increased by approximately 30-fold in infarct macrophages compared with their precursor monocytes (Figure 2J, blue arrow), indicating strong induction by the ischemic tissue environment. Interestingly, even though prior reports have demonstrated the role of IL1B in aggravating post-MI remodeling and heart failure^28–31^, IL1B generated by macrophages in inducing inflammatory injury after MI is an area with surprisingly very limited (if any) data. These data suggest that mDAMP-triggered inflammatory signaling is likely TLR-driven and plays a dominant role in macrophage activation during acute myocardial ischemia, thus inhibitors of macropinocytosis that downregulate this signaling could dampen the inflammatory response post-MI and protect the heart from ischemic injury.

### EIPA Mitigates TLR ligand-Induced Cardiac Inflammation In Vivo

Because mDAMP elicits inflammatory signaling similar to canonical TLR4 activation, including coordinated induction of NF-κB, ERK, mTORC1, and robust *Il1b* expression, and because both EIPA and NPS markedly suppressed mDAMP-induced *Il1b*, we hypothesized that TLR4 signaling may represent a principal pathway sensitive to macropinocytosis inhibition. To directly test this possibility, we examined whether macropinocytosis inhibitors attenuate macrophage responses to the canonical TLR4 ligand lipopolysaccharide (LPS). BMDMs were stimulated with LPS (10 ng/ml) in the presence of vehicle, EIPA, or NPS. Both EIPA and NPS markedly blunted LPS-induced expression of *Il1b* and *Cxcl10* (Figure 3A). EIPA reduced IL1B protein production after LPS stimulation (Figures 3B, 3C). Furthermore, EIPA suppressed LPS-induced interferon-stimulated gene (ISG) expression, including *Cxcl10* (Figure 3D), *Ifit1*, *Ifit3*, *Isg15*, *and Rsad2* (Figure 3E). Collectively, these data support a model in which macropinocytosis-dependent processes are required for maximal macrophage responses to both damage-associated molecular patterns and LPS, and that pharmacologic inhibition of macropinocytosis antagonizes TLR4-driven inflammatory signaling.

**Figure 3.**
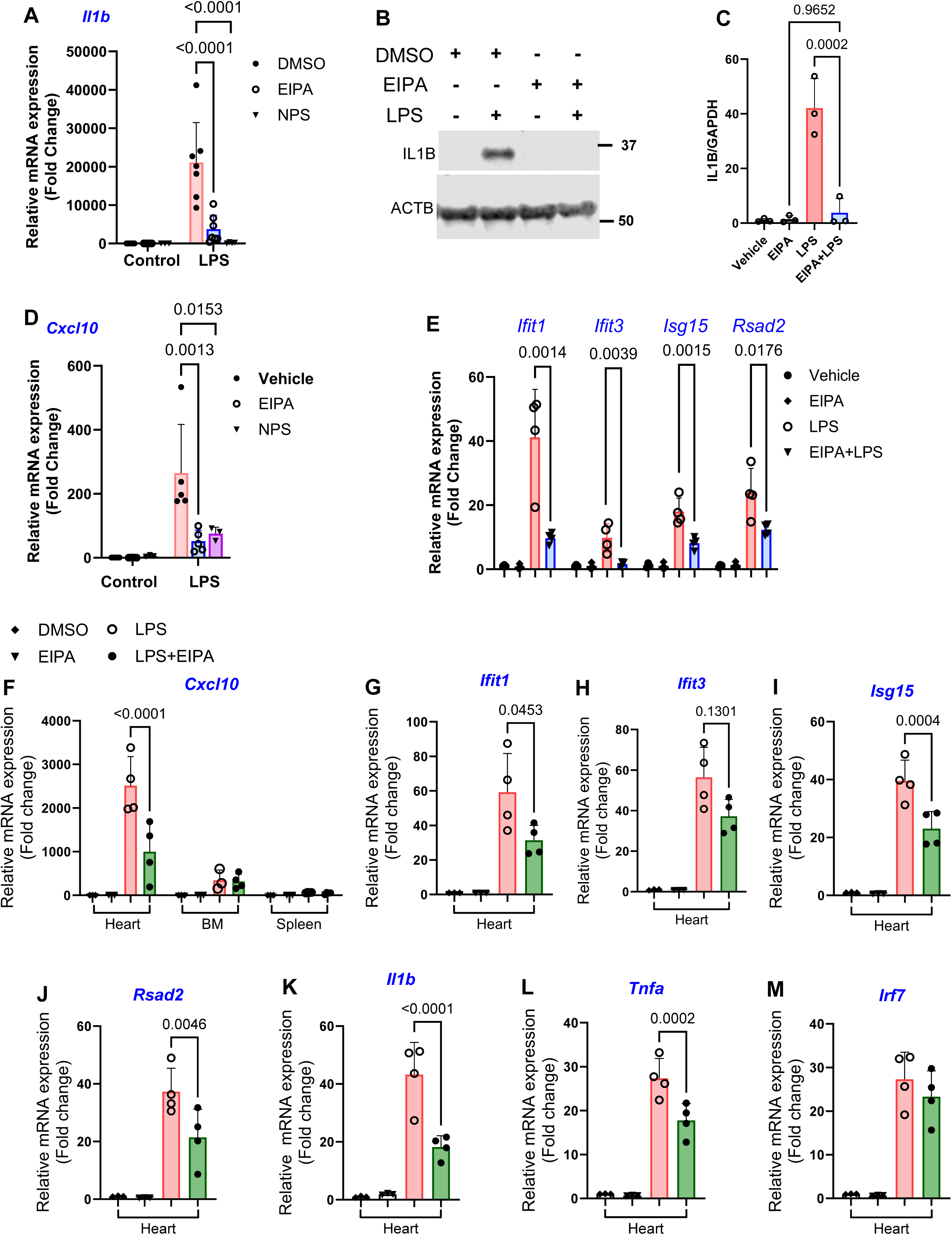
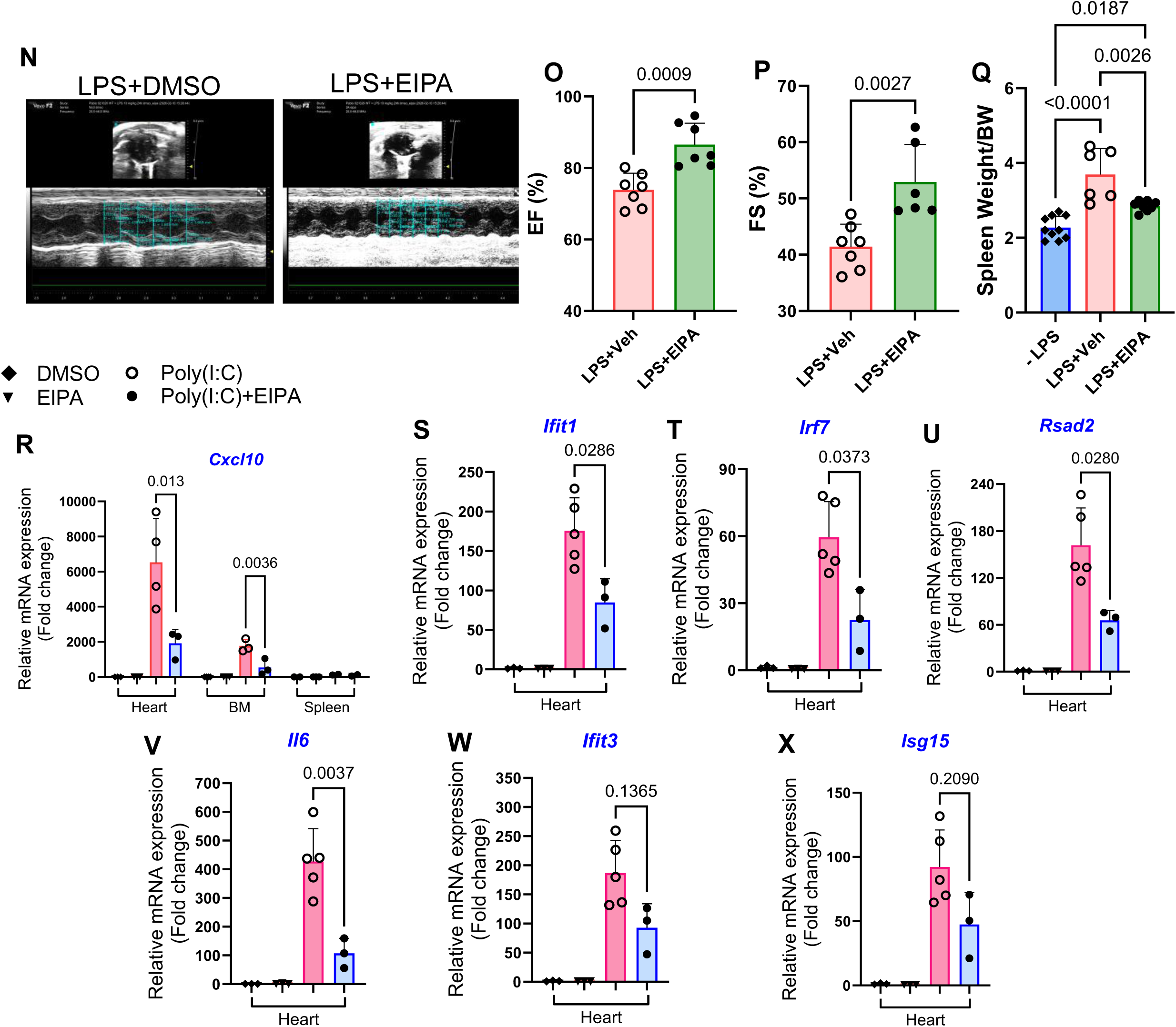
Pharmacologic SLC9A inhibition attenuates inflammatory responses to systemic LPS and poly(I:C) challenge. Because mDAMP induces signaling features consistent with Toll-like receptor activation (Supplementary Figures), we next tested whether macrophage responses to the defined ligands LPS and poly(I:C) are similarly sensitive to macropinocytosis inhibition.A. BMDMs were stimulated with LPS (10 ng/mL) for 4 h in the presence or absence of EIPA or NPS. *Il1b* expression was measured by RT-qPCR. B-C. Immunoblot analysis of IL1B protein expression in BMDMs stimulated with LPS with or without EIPA. D-E. Expression of interferon-stimulated genes (ISGs) in BMDMs following LPS stimulation in the presence or absence of EIPA, including *Cxcl10*, *Ifit1*, *Ifit3*, *Isg15*, and *Rsad2*. F-M. C57BL/6J mice were treated with EIPA (two doses administered before LPS challenge) and injected intraperitoneally with LPS (1 mg/kg). Four hours after LPS administration, heart, spleen, and bone marrow (BM) were harvested for RNA extraction and RT-qPCR analysis. Relative inflammatory responses across tissues are shown using Cxcl10, which is the most robust up-regulated gene. (F). Expression of *Ifit1*, *Ifit3*, *Isg15*, *Rsad2*, *Il1b*, and *Tnfa* in the heart was reduced by EIPA treatment (G-L), whereas Irf7 expression was not significantly altered (M). N-Q. Mice were challenged with LPS (10 mg/kg) after pretreated with daily DMSO or EIPA (12.5mg/kg) for three days. Echocardiography was performed at 6 h after LPS administration. Representative echocardiographic images and quantification of ejection fraction (EF) and fractional shortening (FS) are shown (N-P). Spleen weights were measured 24 h after LPS injection (Q). R-X. Mice were challenged with Poly(I:C) (3 mg/kg). Tissues were collected for RT-qPCR analysis. Relative responses across heart, spleen, and BM are shown (R). Cardiac expression of *Cxcl10*, *Ifit1*, i*rf7*, *Rsad2*, and *Il6* was reduced by EIPA treatment (S-V). Changes in *Ifit3* and *Isg15* did not reach statistical significance (W-X).Statistical analysis: Two-way Anova and Student’s t-test.

To determine whether these effects extend in vivo, we next assessed the impact of EIPA on LPS-induced systemic and cardiac inflammation. This experiment has translational relevance, as EIPA is an amiloride derivative related to clinically available Na+/H+ exchange inhibitors. Mice were challenged systemically with LPS (1 mg/kg) and co-treated with vehicle or EIPA. Tissues were harvested 4 hours after LPS administration for RT-qPCR analysis. Surprisingly, the heart exhibited a more robust inflammatory response to LPS than other immune cell-rich organs such as bone marrow and spleen (Figure 3F). EIPA treatment significantly attenuated LPS-induced cardiac inflammation, reducing expression of ISGs (*Cxcl10*, *Ifit1*, *Ifit3*, *Isg15*, *Rsad2*) and pro-inflammatory cytokines (*Il1b*, *Tnfa*), while exerting minimal effects on *Irf7* (Figures 3F-3M). To assess functional consequences, we employed a higher LPS dose (10 mg/kg) that was sufficient to induce acute systemic inflammation and cardiac dysfunction. Echocardiographic assessment 6 hours after LPS administration revealed reduced ejection fraction (EF) and fractional shortening (FS) (Figures 3 N-3P). EIPA treatment preserved cardiac function and prevented LPS-induced systolic impairment. Additionally, EIPA reduced LPS-induced splenomegaly (Figure 3Q), consistent with attenuation of systemic inflammatory activation.

To determine if EIPA prevents inflammation triggered by other TLRs, we next challenged mice with Poly(I:C), a classical ligand for TLR3. Again, we the heart had the most robust inflammatory response to Poly(I:C) among the spleen and BM (Figure 3R). EIPA treatment attenuated Poly(I:C)-induced ISG and cytokine expression, including *Cxcl10*, *Ifit1*, *Irf7,* and *Il6*.

Collectively, these findings demonstrate that pharmacologic inhibitors of macropinocytosis not only suppressed TLR4-driven inflammatory signaling in macrophages but also mitigated cardiac inflammation and acute cardiac dysfunction induced by TLR activation.

### Deleting *Slc9a1* in monocytes and monocytes-derived macrophages

EIPA is not a specific SLC9A1 inhibitor even though its potent inhibition of macropinocytosis is attributed to SLC9A1/NHE1 inhibition^15, 16^. On the other hand, recent genetic studies indicate that SLC9A1 is not required for macropinocytosis in all cell types^32^. Therefore, we determined the SLC9A isoform expression in bone marrow (BM) monocytes. RNA sequencing of purified monocytes (CD45^+^CD11b^+^Ly6C^hi^Ly6G^-^CD3^-^) showed that *Slc9a1* is the main isoform expressed by monocytes (Figure 4A).

**Figure 4.**
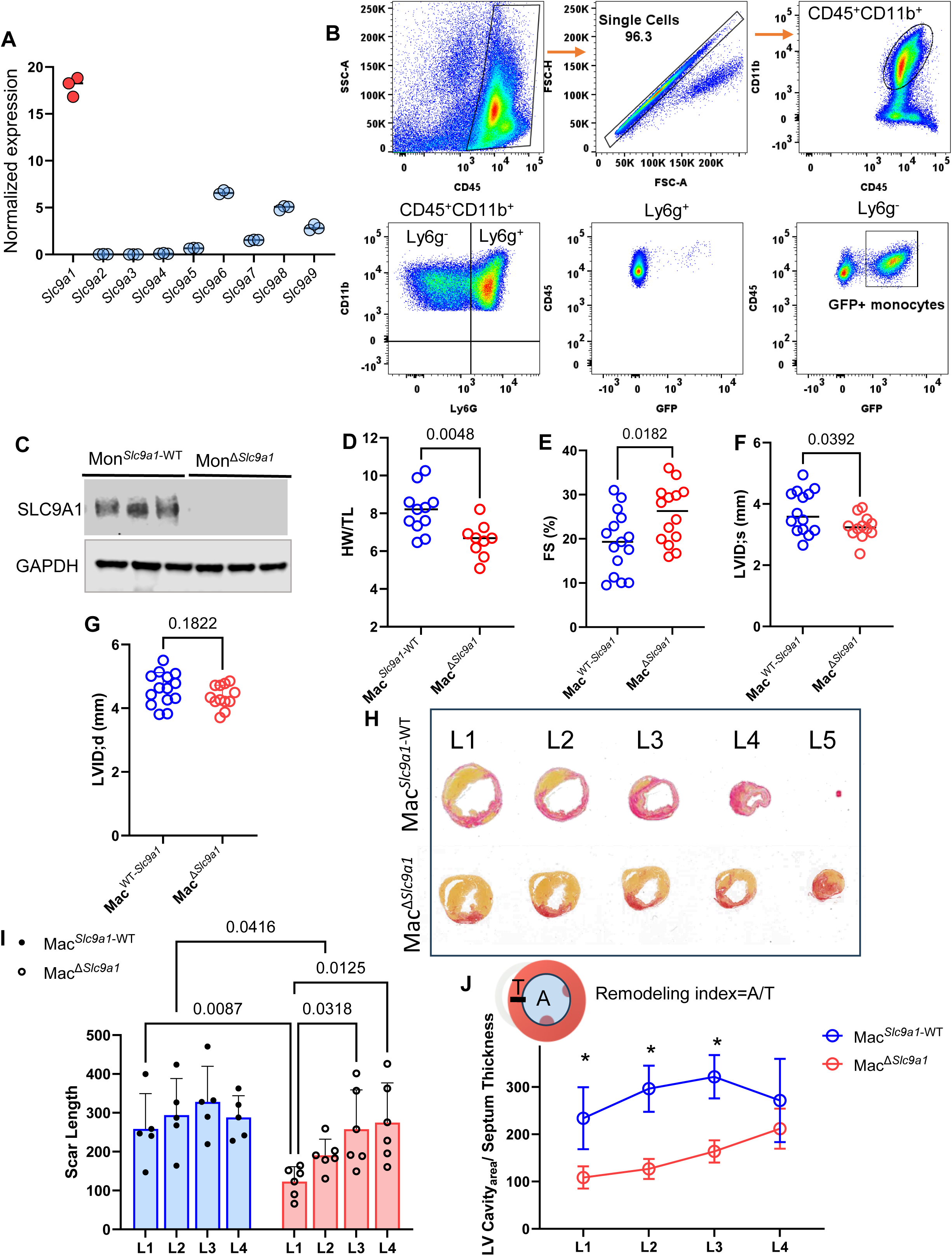
Monocyte- and monocyte-derived macrophage-specific deletion of *Slc9a1* attenuates post–myocardial infarction remodeling. A. Bulk RNA-seq analysis of bone marrow monocytes purified by FACS showing normalized expression of SLC9A family members. *Slc9a1* was the predominant isoform expressed. *Slc9a1*^f/f^;CCR2-Cre-GFP mice and control littermates (*Slc9a1*^f/f^ or CCR2-Cre-GFP) were treated with tamoxifen (75 mg/kg/day, i.p.) for three consecutive days. Mice were subsequently designated as <u>mon</u>ocyte-specific knockout (Mon)^Δ*Slc9a1*^ and control (Mon*^Slc9a1-^*^WT^). B. Gating strategy for isolating bone marrow monocytes expressing endogenous GFP. C. Immunoblot analysis of SLC9A1 protein in bone marrow monocytes purified from Mon^Δ*Slc9a1*^ and CCR2-Cre-GFP mice, confirming efficient deletion. D-F. Myocardial infarction was induced by LAD ligation in Mon^Δ*Slc9a1*^ and Mon*^Slc9a1-^*^WT^ mice. Cardiac remodeling parameters were assessed 7 dpi. Monocyte-derived macrophage-specific deletion of Slc9a1 (Mac^Δ*Slc9a1*^) reduced heart weight/tibia length ratio (HW/TL) (D), improved fractional shortening (FS) (E), decreased left ventricular end-systolic dimension (F), while having no effect on diastolic dimensions (G). H. Representative heart cross-sections from base to apex illustrating left ventricular morphology and infarct scar. I. Quantification of scar length from serial sections corresponding to the levels shown in (G). J. Remodeling index (RI), calculated as the ratio of left ventricular diameter to septal thickness (diagram shown), in Mac^Δ*Slc9a1*^ and Mac*^Slc9a1-^*^WT^ mice. Student’s t-test.

Since BM monocytes are the precursors of macrophages in the infarct tissue, we therefore generated a compound mouse model that carried alleles of Flox *Slc9a1*^fl/fl^ and the CCR2-CreER-GFP. *Slc9a1*^fl/fl^ only and CCR2-CreER-GFP littermates were used as controls. We administered tamoxifen (TMX) at 75 mg/kg body weight for three days to induce *Slc9a1* deletion in monocytes, and the control littermate received the same treatment. After TMX injection, we annotated the *Slc9a1*^fl/fl^;CCR2-CreER-GFP mice as <u>Mon</u>ocyte(Mon)^ΔSlc9a1^ and *Slc9a1*^fl/fl^-only or CCR2-CreER-GFP-only mice as

Mon*^Slc9a1-^*^WT^. The nomenclature of Mon*^Slc9a1-^*^WT^ and Mon^Δ*Slc9a1*^ refers to monocytes from the BM, while <u>mac</u>rophage(Mac)^Slc9a-WT^ and Mac^Δ*Slc9a1*^ are for post-MI macrophages from the cardiac tissue. To confirm SLC9A1 deletion specifically in monocytes, we purified monocytes (CD45^+^CD11b^+^Ly6G^-^GFP^+^) from the BM and performed immunoblot using protein samples extracted from the purified cells. Figure 4B depicts the gating strategy for isolating GFP+ monocytes. Figure 4C shows the loss of SLC9A1 in GFP^+^ monocytes isolated from Mon^Δ*Slc9a1*^ mice compared to the GFP^+^ monocytes from CCR2-CreER-GFP only mice.

### Macrophage-specific *Slc9a1* deletion protects the heart from acute ischemia

To determine if deleting *Slc9a1* in MI macrophages recapitulates the cardiac protection from EIPA, we generated myocardial infarction model by performing LAD ligation in Mac ^Δ*Slc9a1*^ and Mac*^Slc9a1-^*^WT^ mouse cohorts. As MI macrophages are exclusively derived from circulating CCR2^+^ monocytes, using the CCR2-Cre conditionally deletes *Slc9a1* in monocyte-derived macrophages (MDM) including those that localize in MI tissue. One week after the MI procedure (7 dpi), we measured cardiac function, left ventricular dimensions, and the gravimetric parameter HW/TL as markers of ischemia-induced cardiac remodeling. Figure 4D demonstrates that deleting *Slc9a1* in MDMs decreased cardiac mass as indicated by lower HW/TL compared to the control mice. We also observed better cardiac function (fractional shortening, FS) (Figure 4E) and smaller left ventricle systolic dimensions (LVID;s) in the Mac^ΔSlc9a1^ cohort (Figure 4F). Figure 4H shows representative serial cross-sections illustrating the extent of scar formation following MI in Mac^Δ*Slc9a1*^ and Mac*^Slc9a1-^*^WT^ mice. Scar length was quantified as a reliable indicator of LV remodeling after MI^1, 11^, and a remodeling index (RI) was calculated as the ratio of LV internal diameter to septal thickness. As summarized in Figures 4I and 4J, monocyte-derived macrophage (MDM)–specific deletion of *Slc9a1* significantly reduced scar size and decreased the RI. Collectively, these findings indicate that SLC9A1 in MDMs promotes adverse post-MI remodeling and contributes to impaired cardiac function.

To determine whether macrophage-specific deletion of *Slc9a1* alters myeloid cell dynamics following MI, we performed flow cytometric analysis of infarct tissue at 3- and 7-dpi. As illustrated in Figures 5A-5B, we profiled four principal myeloid populations: (1) total macrophages (CD45 ⁺CD11b⁺Ly6G⁻); (2) GFP⁺ MDMs (CD45⁺CD11b⁺Ly6G⁻GFP⁺); (3) recently recruited inflammatory monocytes (CD45⁺CD11b⁺Ly6G⁻GFP⁺Ly6C^hi^); and (4) neutrophils (CD45⁺CD11b⁺Ly6G⁺Ly6C^int^GFP⁻). At 3-dpi, deletion of *Slc9a1* resulted in a significant reduction in total macrophages, GFP⁺ MDMs, and newly recruited Ly6C^hi^ inflammatory monocytes within the infarcted myocardium (Figures 5E-5G). In contrast, neutrophil numbers were modestly increased in Mac^Δ*Slc9a1*^ mice at this early time point (Figure 5H). By 7-dpi, however, the numbers of neutrophils, Ly6C^hi^ monocytes, and GFP⁺ MDMs were comparable between control and knockout cohorts (Figures 5I-5L).In summary, conditional deletion of *Slc9a1* transiently impaired early monocyte recruitment and MDM accumulation following MI, particularly during the acute inflammatory phase, but did not produce sustained alterations in myeloid cell composition at later stages of remodeling.

**Figure 5.**
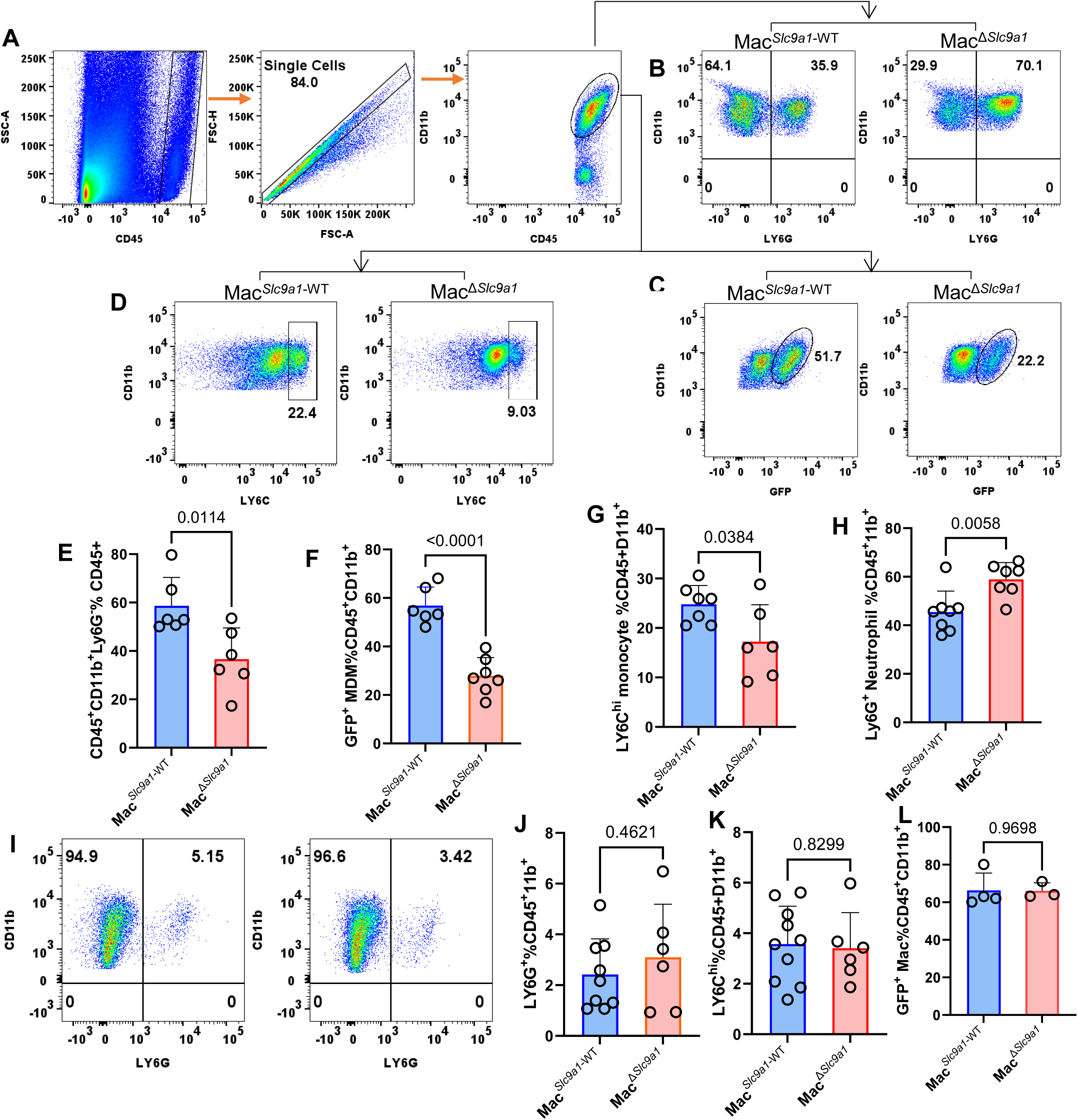
Immune cell profiling in Mac^Δ*Slc9a1*^ and Mac*^Slc9a1-^*^WT^ mice. after MI. A-D. Gating strategy for analysis of myeloid cells from single-cell suspensions prepared from infarct tissue. Cells were first gated as CD45⁺CD11b⁺, which is further separated into the following myeloid subpopulations: total macrophages (CD45⁺CD11b⁺Ly6G⁻) (B), monocyte-derived macrophages (MDMs; CD45⁺CD11b⁺Ly6G⁻GFP⁺) (C), and recently recruited monocytes (CD45⁺CD11b⁺Ly6G⁻Ly6C^hi^) (D). Neutrophils were defined as CD45⁺CD11b⁺Ly6G⁺Ly6C^int^GFP⁻. E-G. Quantification of total macrophages (E), MDMs (F), and newly recruited monocytes (G) in infarct tissue at 3dpi. H. Quantification of neutrophils in infarct tissue at 3dpi. I-J. Neutrophil abundance in infarct tissue at 7dpi. K-L. Quantification of monocytes and MDMs in infarct tissue at 7dpi. Student’s t-test.

### SLC9A1 promotes type I interferon pathway activation in monocyte-derived macrophages of the MI

To elucidate the mechanisms by which SLC9A1 regulates cardiac MDM function following MI, we purified myeloid cells (CD45⁺CD11b⁺) from infarcted hearts at 4-dpi and subjected them to single-cell RNA sequencing using the 10x Genomics platform. Unsupervised clustering identified eight distinct cell populations, including four macrophage clusters (Mac1 -Mac4), neutrophils, dendritic cell-like (DC-like) cells, a small population of Mki67⁺ proliferating cells, and rare lymphocytes (Figure 6A). Canonical marker genes for each cluster are shown in Figure 6B. We focused subsequent analyses on the macrophage populations.

**Figure 6.**
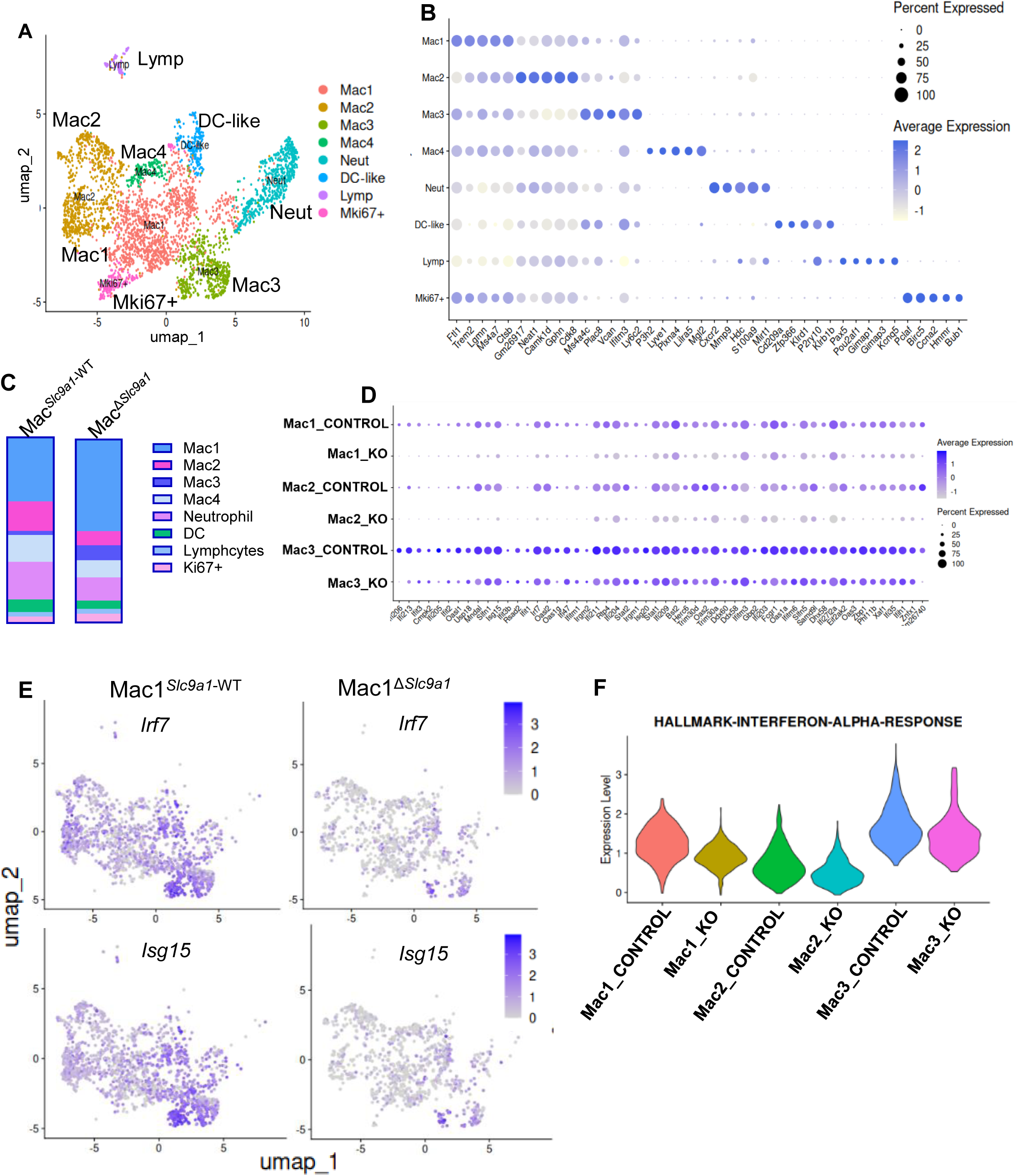
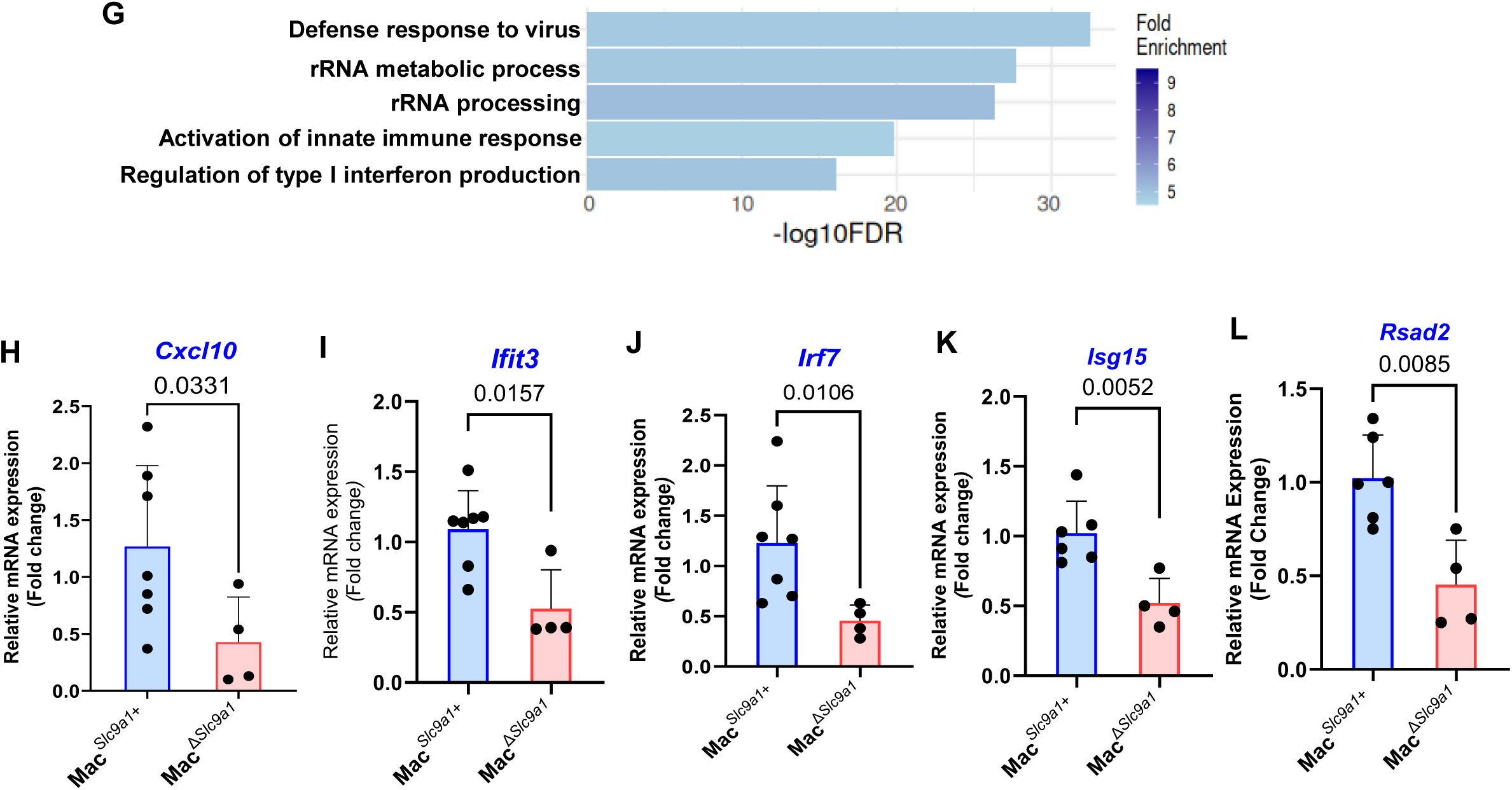
SLC9A1 promoted interferon stimulated genes response in acute ischemia in MDM. Myeloid cells were isolated from infarct tissue 4dpi. CD45⁺ cells were first enriched using magnetic bead separation and subsequently purified by FACS as CD45⁺CD11b⁺CD3⁻ cells. Purified cells were subjected to single-cell RNA sequencing using the 10x Genomics platform. A. UMAP projection showing eight cell clusters identified in the dataset. B. Marker genes used for cluster annotation. C. Proportion of cells in each cluster. D. Dot plot showing the average expression of 48 ISGs across macrophage clusters (Mac1, Mac2, and Mac3) in control and Mac^Δ*Slc9a1*^ samples. E. UMAP feature plots showing expression of *Irf7* and *Isg15*. F. Violin plots showing enrichment scores for the Hallmark Interferon Alpha Response gene set in macrophage clusters (Mac1-Mac3) from control and Mac^Δ*Slc9a1*^ infarct hearts. G. Gene Ontology (GO) enrichment analysis of pathways downregulated in Mac1 cells from Mac^Δ*Slc9a1*^ mice. H-L. RT-qPCR validation of ISG expression in infarct tissue collected 3dpi, including *Cxcl10* (H), *Ifit3* (I), *Irf7* (J), *Isg15* (K), and *Rsad2* (L). Student’s t-test.

The *Flt1*⁺*Trem2*⁺*Lgmn*⁺*Ms4a7*⁺*Ctsb*⁺ cluster (Mac1) exhibited a transcriptional signature consistent with reparative, efferocytosis-enriched macrophages. Genes such as *Lgmn*, *Trem2*, and *Ctsb* are associated with phagocytic clearance and resolution of inflammation ^33, 34^, while *Trem2* also mediates lipid sensing and resolution programs^34, 35^. Expression of *Flt1* (*Vegfr1*) suggests potential involvement in angiogenic or VEGF-related signaling. The enrichment of efferocytosis-associated and pro-resolution pathways indicates that Mac1 likely represents emerging reparative, M2 -like macrophages that arise during the second half of the first week following MI. Consistent with this interpretation, *Ms4a7* has been implicated in promoting M2-like polarization and immune suppression^36^.

The *Gm26917*⁺*Neat1*⁺*Cdk8*⁺*Camk1d*⁺ cluster (Mac2) displayed a transcriptional profile indicative of interferon and NFKB pathway activation. Both *Neat1* and *Gm26917* are long noncoding RNAs associated with inflammasome activation and TLR/NFKB–mediated inflammatory responses^36, 37^. CDK8 encodes a transcriptional kinase that amplifies IFN- and NFKB-dependent gene programs, while *Camk1d* suggests Ca²⁺-dependent pro-inflammatory signaling^38, 39^. Together, these features define Mac2 as a strongly pro-inflammatory macrophage population.

The *Ms4a4c*⁺*Plac8*⁺*Vcan*⁺*Ifitm3*⁺*Ly6c2*⁺ cluster (Mac3) likely represents recently recruited monocyte-derived macrophages in the injured heart. High expression of *Ly6c2* and *Plac8* reflects monocyte-associated signatures^40^, while *Vcan* indicates engagement with extracellular matrix and vascular interfaces. The presence of the interferon-stimulated gene *Ifitm3* further supports an activated, newly infiltrating phenotype. This profile is consistent with prior single-cell studies of muscle and nerve injury in which Plac8 and Ifitm3 marked infiltrating and activated monocyte–macrophage subsets^41^.

Finally, the Mac4 cluster was characterized by expression of *Lyve1*, a marker of tissue-resident macrophages, and enrichment of *Mgl2* (*CD301b*), commonly associated with resident, M2-like macrophages^42^. These features suggest that Mac4 represents cardiac resident macrophages. The presence of this population likely reflects inclusion of border-zone myocardium during cell isolation, despite efforts to enrich primarily for infarct tissue. The proportion of each cluster is depicted in Figure 5C.

### SLC9A1 promotes interferon stimulated gene response in MI

Using the scRNA-seq dataset, we performed differentially expressed gene (DEG) analysis comparing the main macrophage clusters (Mac1,2,3) between Mac^Δ*Slc9a1*^ and Mac*^Slc9a1-^*^WT^ mice. Strikingly, interferon-stimulated genes (ISGs) emerged as the predominant transcriptional program downregulated upon deletion of *Slc9a1*. Among the top 20 most significantly downregulated genes, approximately 80% were ISGs. Figure 6D illustrates the expression of 46 ISGs identified within the top 100 differentially expressed genes across the Mac1, Mac2, and Mac3 clusters. Deletion of *Slc9a1* resulted in broad suppression of these ISGs across macrophage subsets. Notably, macrophages within the Mac3 cluster exhibited the highest baseline expression of ISGs compared to Mac1 and Mac2. This cluster likely corresponds to the previously described IFN cardiac macrophage population^2^. Consistent with this interpretation, Mac3 cells display features of recently recruited monocyte-derived macrophages, including high expression of *Ly6c2*, aligning with prior reports that post-MI ISG responses are initiated in the bone marrow^43^. Representative UMAP heatmaps (Figure 6E) further demonstrate reduced expression of canonical ISGs, including *Isg15* and *Irf7*, in *Slc9a1*-deficient macrophages compared with wild-type controls.

Gene Set Enrichment Analysis (GSEA) confirmed these observations. The most significantly enriched pathway among downregulated genes was the Hallmark IFNA response signature (Figure 6F). Consistently, Gene Ontology (GO) analysis revealed enrichment of pathways related to defense response to virus, type I interferon production, and innate immune activation (Figure 6G), supporting a central role for SLC9A1 in regulating interferon-driven inflammatory programs.

To independently validate the reduction in type I interferon signaling, we performed quantitative RT - PCR on RNA isolated from infarct tissue of Mac^Δ*Slc9a1*^ and Mac*^Slc9a1^*^-WT^ mice. As shown in Figures 6H-6L, macrophage-specific deletion of *Slc9a1* significantly reduced expression of key ISGs, including *Cxcl10*, *Ifit3*, *Irf7*, *Isg15*, and *Rsad2*. These findings corroborate the single-cell RNA-sequencing results and establish SLC9A1 as a critical promoter of type I interferon pathway activation in infarct - associated macrophages.

### SLC9A1 Enhances Interferon Responses by Facilitating Poly(I:C) Signaling

To define the mechanism by which SLC9A1 regulates type I interferon (IFN) and ISG responses, we focused on the principal endosome-dependent signaling pathways linking endocytosis to IFN induction - namely TLR3 and TLR4. TLR3 recognizes double-stranded RNA mimetics such as Poly(I:C), whereas TLR4 is activated by lipopolysaccharide (LPS). Engagement of these receptors triggers NFKB activation and, following ligand or ligand-receptor internalization, IRF3-dependent signaling that drives expression of ISGs and inflammatory cytokines (e.g., *Il1b*, *Il6*)^44^.

To directly assess the role of SLC9A1, we performed siRNA-mediated knockdown in BMDMs. Efficient reduction of SLC9A1 protein levels was confirmed by immunoblotting (Figure 7A). Macrophages were then stimulated with free Poly(I:C) (without transfection reagents) to selectively engage endosomal TLR3 signaling. As shown in Figures 7B–7F, *Slc9a1* knockdown significantly reduced Poly(I:C)-induced expression of ISGs, including *Cxcl10*, *Ifit1*, and *Isg15*, as well as inflammatory cytokines *Il1b* and *Il6*. In contrast, *Irf7* expression was not significantly affected (Figure 7G). These findings indicate that SLC9A1 facilitates TLR3-dependent inflammatory and ISG responses.

**Figure 7.**
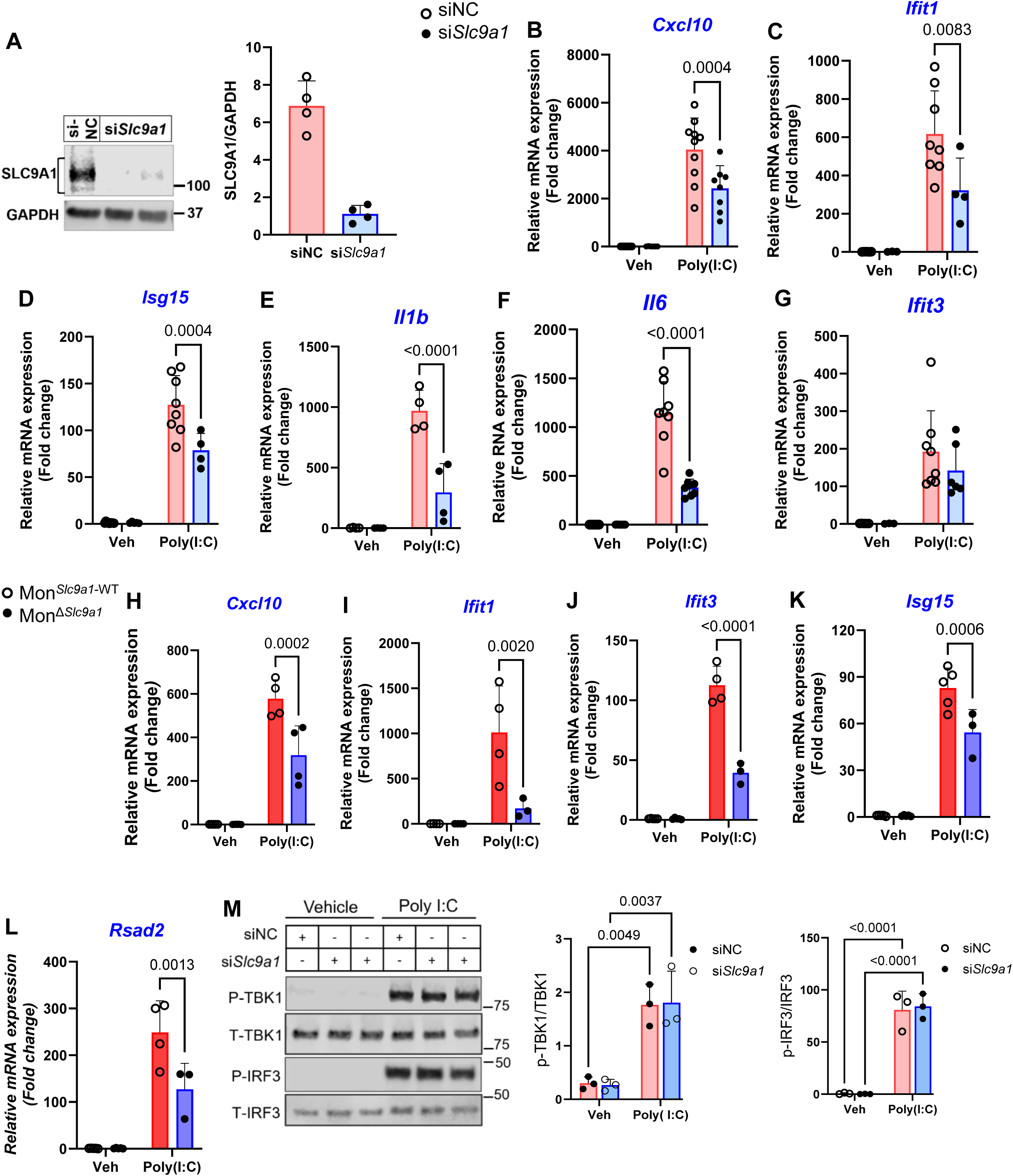
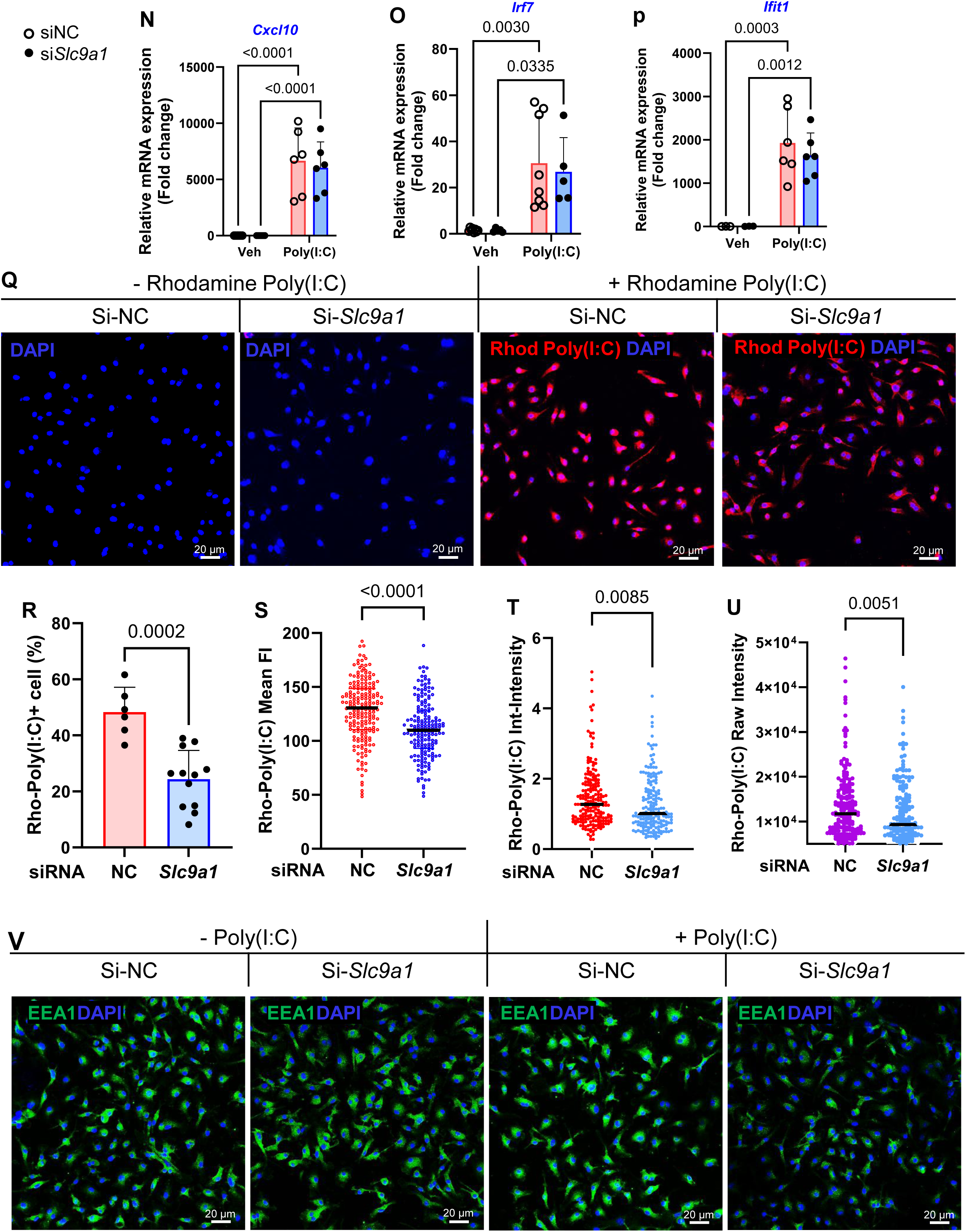
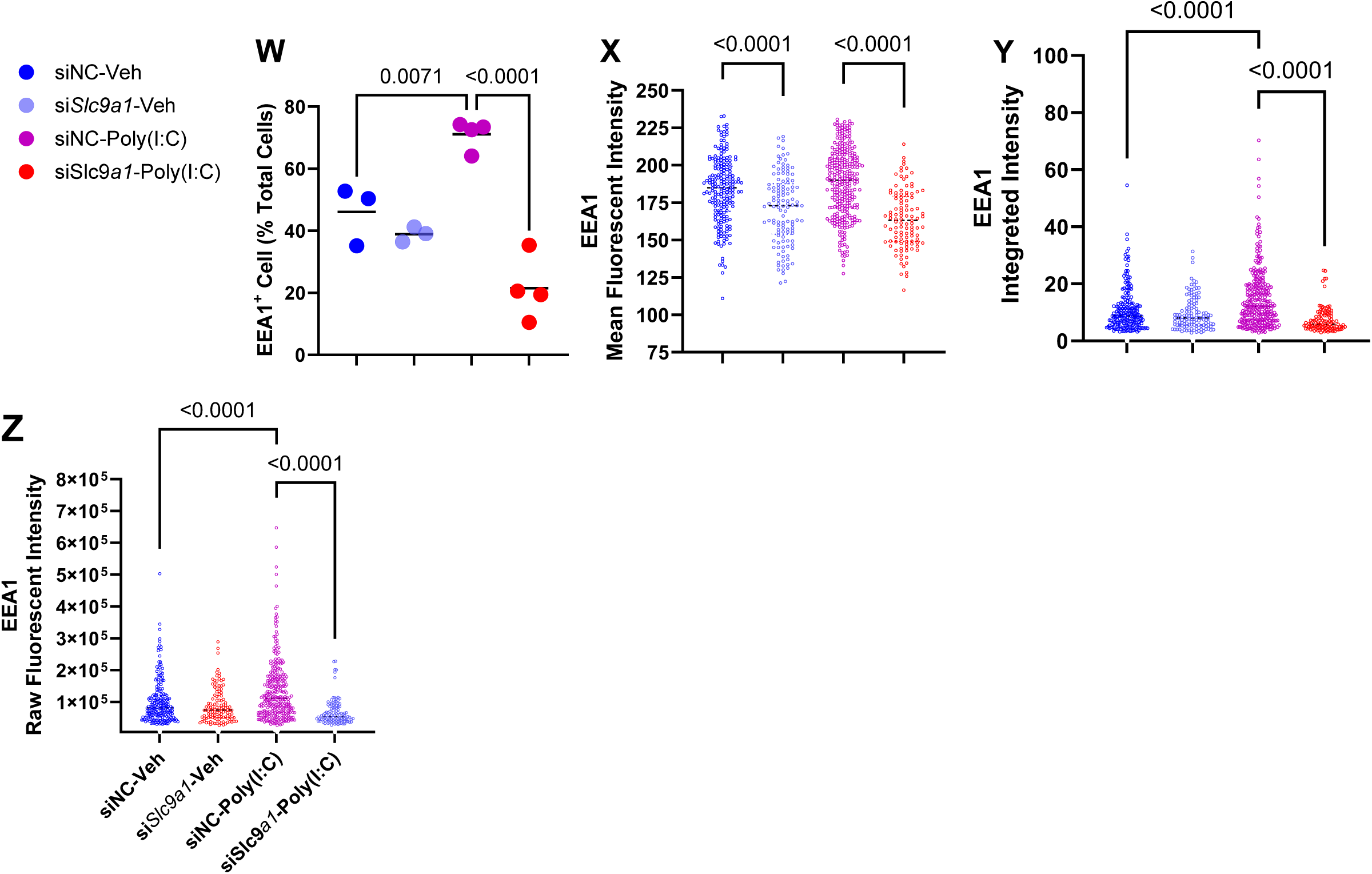
SLC9A1 regulates poly(I:C)-induced signaling and ligand uptake in macrophages. A. BMDMs were treated with control siRNA (siNC) or *Slc9a1* siRNA (si*Slc9a1*). SLC9A1 protein levels were assessed by immunoblot analysis four days after siRNA treatment. B-F. BMDMs were stimulated with poly(I:C) at 5 μg/ml for 4 h after siRNA treatment. Expression of ISGs and inflammatory cytokines was measured by RT-qPCR, including *Cxcl10* (B), *Ifit1* (C), *Isg15* (D), *Il1b* (E), and *Il6* (F). G. Expression of *Ifit3* following poly(I:C) stimulation in siNC- and si*Slc9a1*-treated BMDMs. H-L. Control mice (Mon*^Slc9a1^*^-WT^) and mice with monocyte-specific deletion of Slc9a1 (Mon^Δ*Slc9a1*^) were generated by tamoxifen administration. The mouse cohorts generated were challenged with Poly(I:C) injection via i.p. at 4 mg/kg x1. Bone marrow monocytes were isolated and ISG expression was quantified by RT-qPCR, including *Cxcl10* (H), *Ifit1* (I), *Ifit3* (J), *Isg15* (K), and *Rsad2* (L). M-P. BMDMs prepared as in (A) were transfected with poly(I:C) (500 ng/mL). Protein and RNA were collected 4 h after transfection. Immunoblot analysis of phosphorylated and total TBK1 and IRF3 is shown and the phosphorylated isoforms were quantified (M). Expression of *Cxcl10* (N), *Irf7* (O), and *Ifit1* (P) was measured by RT-qPCR. Q. BMDMs treated as in (A) were incubated with rhodamine-labeled poly(I:C) for 1 h. Representative confocal images of ligand uptake are shown. R-U. Quantification of rhodamine-Poly(I:C) uptake using ImageJ, including the percentage of cells with detectable signal (R), mean fluorescence intensity (S), integrated intensity (T), and raw fluorescence intensity (U). V. BMDMs were stimulated with Poly(I:C) (5 μg/ml) for 4 h and stained with the early endosome marker EEA1. Representative confocal images are shown. W-Z. Quantification of EEA1-positive endosomes using ImageJ, including the percentage of cells with detectable endosomes with a set threshold (W), mean fluorescence intensity (X), integrated intensity (Y), and raw fluorescence intensity (Z). Student’s t-test and two-way Anova.

To extend these observations in vivo, we administered Poly(I:C) intraperitoneally (4 mg/kg body weight) to Mon^Δ*Slc9a1*^ mice and control littermate controls (Mon*^Slc9a1-^*^WT^). Bone marrow monocytes (CD45⁺CD11b⁺GFP⁺) were isolated four hours after injection to assess activation of the TLR3 -IFN-ISG axis. As shown in Figures 7H-7L, *Slc9a1*-deficient monocytes exhibited significantly attenuated induction of *Cxcl10*, *Ifit1*, *Ifit3*, *Isg15*, and *Rsad2* compared with controls. Collectively, these in vitro and in vivo data demonstrate that SLC9A1 promotes TLR3-mediated interferon signaling and supports a broader role for SLC9A1 in regulating endosome-dependent innate immune pathways.

### SLC9A1 Has Limited Impact on Cytosolic Nucleic acid Receptor-Mediated IFN–ISG Activation

In addition to endosomal TLR3, double-stranded RNA (dsRNA) such as Poly(I:C) can be detected in the cytosol by the RNA sensors IFIH1 (MDA5) and RIG-I, leading to robust activation of the TBK1–IRF3 axis and subsequent induction of type I interferon and ISGs. To determine whether SLC9A1 also regulates IFIH1/RIG-I–dependent signaling, we delivered Poly(I:C) directly into the cytosol by transfection following siRNA-mediated knockdown of Slc9a1 in BMDMs. As expected, transfected Poly(I:C) induced strong phosphorylation of TBK1 and IRF3, confirming activation of the cytosolic RNA-sensing pathway (Figure 7M). In contrast to stimulation with free Poly(I:C), which requires endosomal uptake, knockdown of *Slc9a1* did not significantly attenuate TBK1 or IRF3 phosphorylation in response to cytosolic Poly(I:C). Consistently, expression of representative ISGs, including *Cxcl10*, *Ifit1*, and *Irf7*, was not reduced following Poly(I:C) transfection in *Slc9a1*-deficient macrophages (Figures 7N-7P). These findings indicate that SLC9A1 selectively facilitates endosome-dependent TLR3 signaling but does not substantially regulate cytosolic RNA sensor–mediated IFN–ISG activation.

### SLC9A1 Promotes Poly(I:C) Endocytosis

Extracellular Poly(I:C) is internalized through receptor-mediated endocytic pathways, including uptake via scavenger receptors ^45, 46^. To determine whether SLC9A1 regulates this process, we tracked the internalization of rhodamine-labeled Poly(I:C) (Rho-Poly(I:C)) in BMDMs. As shown in Figure 7Q, Rho-Poly(I:C) was efficiently internalized by control macrophages. In contrast, knockdown of *Slc9a1* reduced intracellular Rho-Poly(I:C) accumulation. Quantitative image analysis using standardized thresholding (ImageJ, see Methods) demonstrated a significant reduction in the proportion of cells exceeding the fluorescence threshold (Figure 7R), as well as decreases in mean fluorescence intensity, integrated intensity, and raw fluorescence intensity in *Slc9a1*-deficient macrophages (Figures 7S-7U).

To further assess endocytic trafficking, we examined early endosome formation by immunostaining for the early endosomal marker EEA1 (Figure 7V). Under basal conditions, no significant differences were observed between control and *Slc9a1*-deficient macrophages in the percentage of EEA1⁺ cells. Poly(I:C) stimulation increased early endosome formation (Figure 7W), and *Slc9a1* knockdown significantly reduced early endosome formation (Figure 7W). Consistently, mean, integrated, and raw EEA1 fluorescence intensities were all diminished in *Slc9a1*-deficient macrophages after Poly(I:C) exposure (Figures 7X-7Z).

Collectively, these results demonstrate that pharmacologic SLC9A inhibition effectively mitigates TLR3- and TLR4-driven cardiac inflammation in vivo. While genetic deletion reveals a selective role for SLC9A1 in endosome-dependent interferon signaling, pharmacologic inhibition exerts broader anti-inflammatory effects, highlighting the therapeutic potential of targeting endocytic-dependent innate immune activation in inflammatory cardiac injury by an existing clinical drug.

## Discussion

In this study, we sought to define the role of macropinocytosis in macrophages during post-MI repair and remodeling. We demonstrated that macropinocytosis is robustly activated in the infarcted heart and that MI-associated macrophages represent the predominant macropinocytic population. Pharmacologic inhibition of macropinocytosis with the amiloride derivative EIPA protected against ischemic injury, improving cardiac function and attenuating adverse remodeling. To define the macrophage-intrinsic mechanisms underlying this protection, we generated monocyte-specific and monocyte-derived macrophage (MDM)-specific deletion of *Slc9a1*, the predominant Na⁺/H⁺ exchanger isoform expressed in these cells and a principal molecular target of EIPA. Conditional deletion of *Slc9a1* phenocopied the cardioprotective effects observed with EIPA treatment, establishing an important role for macrophage SLC9A1 in ischemic injury responses.

Mechanistically, loss of *Slc9a1* markedly suppressed interferon-stimulated gene (ISG) expression in monocytes and MDMs. Using complementary in vitro and in vivo approaches, we demonstrate d that SLC9A1 facilitates TLR3-dependent signaling by promoting ligand endocytosis and delivery to endosomal signaling compartments. Notably, genetic deletion of *Slc9a1* did not fully recapitulate the inhibitory effects of EIPA on TLR4 signaling, indicating that EIPA has other targets beyond SLC9A1

Additionally, EIPA robustly suppressed cardiac inflammatory responses induced by systemic Poly(I:C) and LPS, extending the relevance of our findings beyond sterile ischemic injury to inflammatory states associated with viral and bacterial triggers. These findings reveal a previously unrecognized divergence between genetic SLC9A1 deletion and pharmacologic NHE inhibition in their effects on TLR3 and TLR4 signaling. Pharmacologic inhibition of SLC9A exerts broader anti-inflammatory effects. In summary, these findings suggest that both SLC9A1-specific strategies and global SLC9A inhibition may hold therapeutic potential in myocardial injury, systemic infection, and inflammation-driven cardiac dysfunction.

### Macropinocytosis as a regulator of Inflammatory Sensing

Macrophages are among the most plastic cells, and their phenotypes are decided through an interplay between changes in their transcriptional programs and environmental cues^8, 9^. In the infarcted myocardium, where large quantities of DAMPs are released, macropinocytosis provides an efficient mechanism to internalize and concentrate these signals. As macropinosomes mature and integrate into the endosomal system, they generate signaling-competent compartments that facilitate interactions between DAMPs and PRRs, thereby promoting activation of the NFKB and IRF3 pathways and leading to inflammatory cytokine and chemokine expression that can exacerbate ischemic injury. Although macropinocytosis has largely been studied as a mechanism of immune surveillance and environmental sampling in macrophages^47^, our findings suggest that this pathway can also contribute to pro-inflammatory signaling in the setting of myocardial infarction.

This study also showed that myocardial DAMP preparations closely recapitulate the endogenous ischemic milieu and elicit signaling profiles characteristic of TLR activation, especially TLR4. Inhibition of macropinocytosis attenuated these responses, supporting the concept that macropinocytosis functions as an upstream key upstream signaling input governing DAMP-induced inflammatory amplification. These findings identify macropinocytosis as an essential regulator of TLR-dependent inflammatory signaling in the injured heart.

### Divergent Effects of Genetic SLC9A1 Deletion and Pharmacologic Inhibition

EIPA is widely used as an inhibitor of plasma membrane Na+/H+ exchangers (NHE/SLC9A) and has the highest affinity for SLC9A1 (IC50 0.01–0.02 μM, substantially lower than that of other SLC9A isoforms)^48^. Given that SLC9A1 is the predominant Na+/H+ exchanger expressed in monocytes and monocyte-derived macrophages (MDMs), it has been generally assumed that EIPA-mediated inhibition of macropinocytosis primarily reflects suppression of SLC9A1 activity. However, our data found that the responses to Poly(I:C) and LPS were differently affected by genetic *Slc9a1* knockdown or deletion compared with pharmacologic inhibition by EIPA. The most notable effect of genetic disruption of *Slc9a1* was impaired cellular responses to extracellular Poly(I:C), consistent with reduced uptake and diminished endosomal delivery of nucleic acid ligands, while deletion of *Slc9a1* had a limited impact on LPS-activated signaling (data not shown). Because Poly(I:C)-induced signaling depends on ligand internalization and access to intracellular compartments, even modest reductions in SLC9A1-dependent membrane dynamics or endocytic flux would be expected to selectively attenuate these responses. In contrast, pharmacologic inhibition with EIPA suppressed inflammatory activation induced by both Poly(I:C) and LPS. This suggests that EIPA exerts broader effects on membrane remodeling, intracellular pH regulation, and endocytic trafficking. Such global perturbations may interfere not only with ligand uptake but also with downstream trafficking or signaling events associated with multiple inflammatory pathways. Thus, whereas SLC9A1 appears to selectively facilitate endosome-dependent nucleic acid sensing, traditional pharmacologic methods of inhibiting produces wider suppression of inflammatory activation. This distinction highlights the overlapping yet non-identical phenotypes observed with genetic and pharmacologic perturbations and underscores the value of genetic approaches in defining isoform-specific immune functions.

### SLC9A1 in Endocytic Regulation

The finding that SLC9A1 selectively promotes the ***signaling downstream*** of TLR3 ligands is intriguing as it implies the role of SLC9A1 in receptor-mediated endocytosis, the currently known path that delivers the ligand to the endosomal receptor TLR3^45, 46^. Although there is no strong evidence that SLC9A1 directly regulates canonical receptor-mediated endocytosis machinery (e.g., clathrin, AP2, dynamin, Rab5/7) by binding to or modifying those components, SLC9A1 regulates key biophysical prerequisites of receptor-mediated endocytosis, including submembranous pH^15^ and actin dynamics^13^, thereby shaping membrane remodeling processes membrane curvature and cortical tension required for efficient endocytic uptake^49,50^. In macrophages, where rapid ligand uptake and endosomal signaling are central to innate immune activation, SLC9A1-dependent control of membrane dynamics may therefore regulate the efficiency with which TLR3 ligands, such as extracellular RNA are internalized and delivered to signaling-competent endosomal compartments. Additionally, it is also possible that the TLR3 ligands are internalized via the non-selective fluid phase uptake (macropinocytosis), which eventually deliver their contents to endo-lysosomes. However, how SLC9A1 selectively engages with the TLR3 ligand endocytosis process remains to be elucidated in future studies.

### Translational Implications of SLC9A Inhibition

To assess translational relevance, we evaluated the effects of pharmacologic NHE inhibition in models of systemic innate immune activation. Despite limited isoform specificity, amiloride derivatives markedly attenuated cardiac inflammation induced by systemic Poly(I:C) and LPS. From a clinical perspective, therapeutic benefit may not require strict isoform specificity if modulation of dominant membrane-trafficking pathways is sufficient to suppress maladaptive inflammation.

In our short-duration systemic LPS and Poly(I:C) challenge models, the robust attenuation of cardiac inflammatory responses by EIPA occurred within a timeframe in which substantial recruitment of circulating immune cells to the myocardium is unlikely. Notably, the heart exhibited a more pronounced inflammatory response to these TLR ligands than immune cell-rich organs such as bone marrow and spleen. These observations raise the possibility that EIPA exerts anti-inflammatory and antiviral effects not only through modulation of monocyte-macrophage function but also through direct actions on resident cardiac cell populations, including cardiomyocytes. Given that cardiomyocytes express functional TLRs and contribute to inflammatory signaling during systemic infection, further investigation into the cell-type–specific effects of SLC9A inhibition is warranted. Such studies may be particularly relevant in conditions such as sepsis-induced cardiomyopathy, where excessive innate immune activation within the myocardium contributes to cardiac dysfunction.

### Limitations

Several limitations warrant consideration. First, although we establish a functional link between SLC9A1 and TLR3-dependent signaling, the precise molecular mechanism by which SLC9A1 regulates ligand endocytosis and endosomal signaling remains to be defined. Second, our study focused on the monocyte–macrophage compartment; the potential contributions of SLC9A1 in other cardiac cell types, including cardiomyocytes, were not investigated. Third, while we focused on 7-dpi to assess remodeling, which is a strong predictor of long-term remodeling, longer observation time, including 30-dpi, is planned for future studies. Fourth, mDAMP, as prepared, showed a signature of TLR4 activation. This does not exclude other TLRs, such as TLR3 activation in the MI environment, because nucleotides may have been lost during the preparation of mDAMP.

### Conclusion

In summary, our study identifies macrophage SLC9A1 as a previously unrecognized regulator of innate immune activation in the injured heart. By facilitating endocytic processes required for TLR3-dependent signaling, SLC9A1 amplifies interferon-driven inflammatory responses that contribute to post–myocardial infarction remodeling. The differential effects observed between genetic *Slc9a1* deletion and pharmacologic inhibition underscore the complexity of membrane trafficking and ion exchange in shaping innate immune signaling pathways. Importantly, we demonstrate that EIPA, an amiloride derivative with established clinical use, exerts potent anti - inflammatory effects in the heart in response to TLR3 and TLR4 agonists, revealing unexpected translational potential. Together, these findings expand the conceptual framework linking Na ⁺/H⁺ exchange, membrane dynamics, and endosome-dependent immune signaling, and provide a foundation for future therapeutic strategies targeting SLC9A-dependent pathways to modulate inflammation in myocardial injury and systemic inflammatory states.

## Statements, Declarations, and Acknowledgement

### Ethical standards

*Animal work described in this manuscript has been approved and conducted under the oversight of the UT Southwestern Institutional Animal Care and Use Committee*.

### Funding

This work was supported by VA MERIT Review award I01 BX004562 (X.L., D.J.C) and National Heart, Lung, and Blood Institute of the National Institutes Health awards R01HL145298 (J.W., X.L., D.J.C), P01 HL160488-01A1 (P.P., G.C. X.L., D.J.C), and T32HL125247 (M.C.M).

### Competing interests

The authors have no relevant financial or non-financial interests to disclose.

### Data availability

The RNA-sequencing data will be deposited into a public repository.

### Authors’ contributions

JW and PP contributed equally and performed most of the experiments. YM performed scRNA-sequencing data analysis. GC and MM contributed to some experiments and generated data for Figures 1, 2, and 3. MM also contributed to manuscript writing and editing. XL generated the MI model. DJC oversaw the project, including experimental design, data analysis and interpretation, and manuscript writing.

## Acknowledgement

We sincerely thank Dr. Dandan Sun from the University of Pittsburgh for providing the *Slc9a1*-flox/flox mice.

---

**Supplemental. Figure 1.**
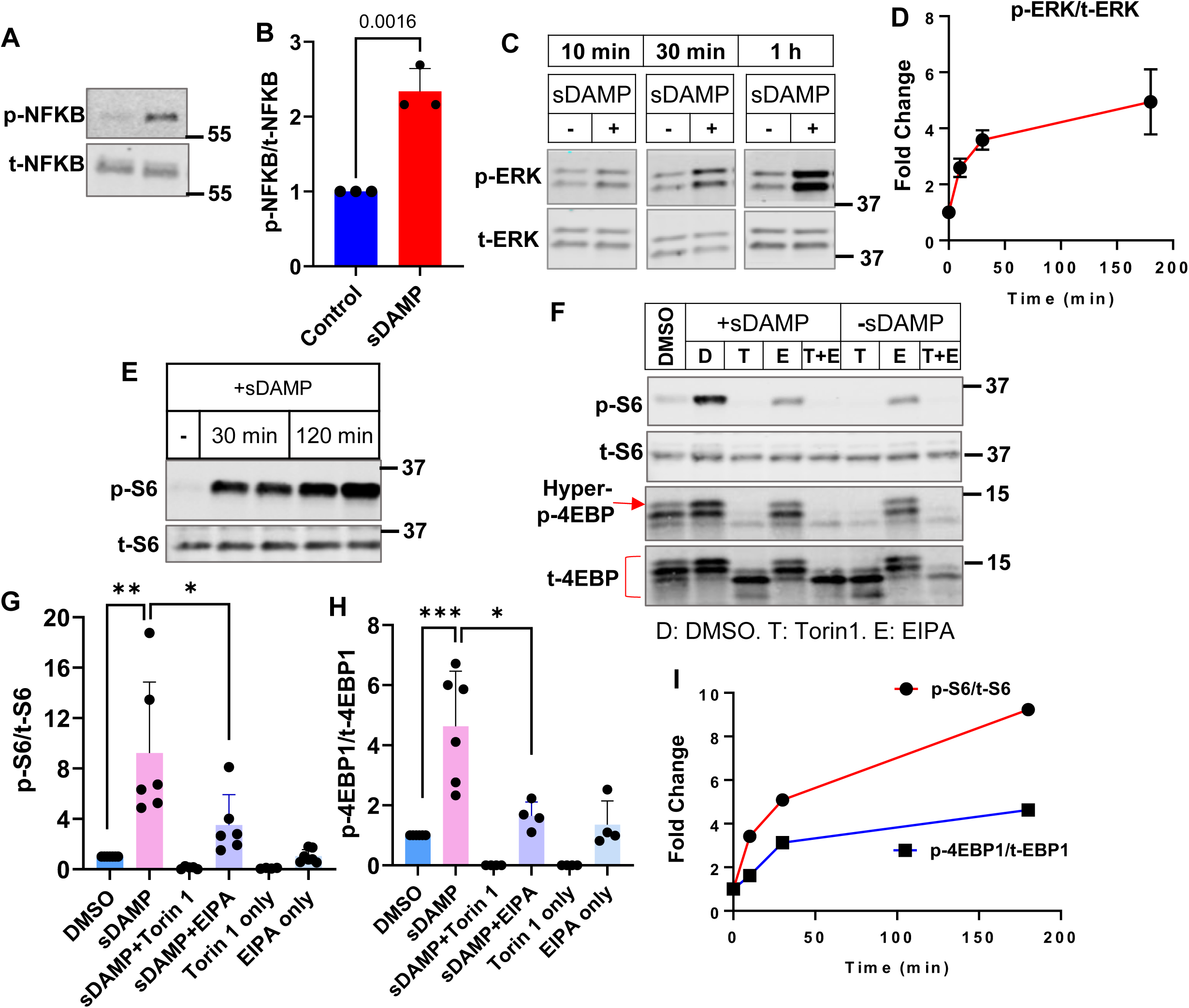
mDAMP simulates TLR agonist by triggering acute activation of NFKB, ERK, and mTORC1. BMDMs were treated with mDAMP. Protein samples extracted after the treatment were subjected to SDS-PAGE and immunoblot analysis. **A and B.** The phosphorylation/activation of NFKB after 4 h treatment of mDAMP. The bar graph depicts summarized results from different experiments. **C and D.** ERK activation by mDAMP. The time course of ERA activation is summarized in D. **E and F.** Phosphorylation of S6 and hyper-phosphorylation of 4EBP1 after 4 h mDAMP stimulation. **G and H.** Summarized results from individual experiments as performed in E and F. **I.** The time course of S6 phosphorylation. Student t-test (B). Two-way Anova (G, H).

**Supplemental. Figure 2.**
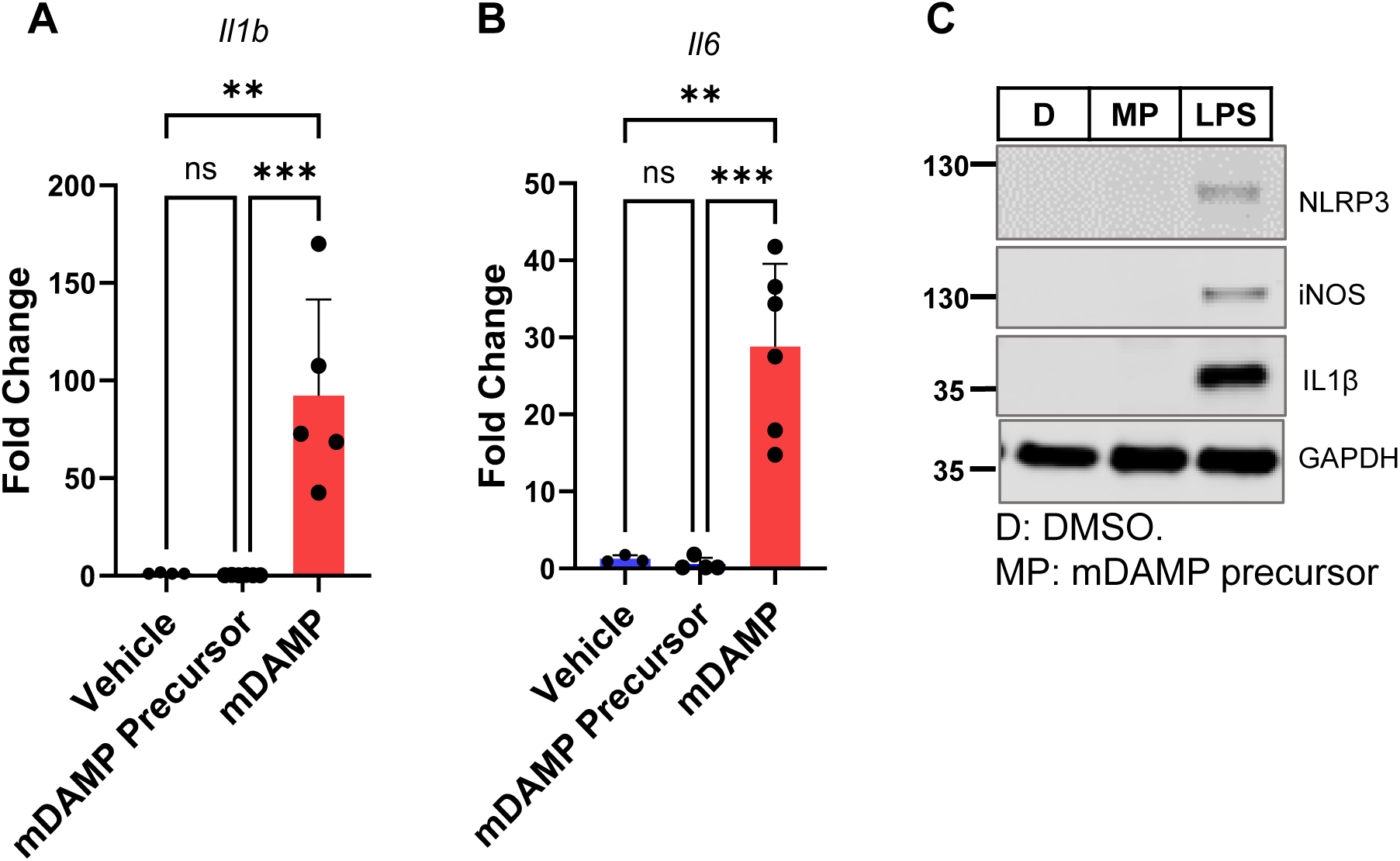
The precursor of mDAMP did not induce inflammatory cytokine expression. Mitochondria were enriched from myocardium obtained from mice at baseline. The preparation is named mDAMP precursor. BMDMs were treated with the precursor or mDAMP. **A and B.** The precursor did not induce inflammatory cytokine expression, including *Il1b* (A) and *Il6* (B), in contrast to mDAMP. **C.** The precursor did not induce NLRP3, INOS, and IL1B protein expression. LPS was used as a positive control. One-way Anova.

## Notes

### Competing Interest Statement

The authors have declared no competing interest.

### Summary of Updates

Revisions were made to improve readability. There are no significant changes in the contents.

